# Heterogeneous causes of acute respiratory distress syndrome correlate with distinct peripheral polyunsaturated fatty acid metabolites

**DOI:** 10.64898/2026.01.09.698377

**Authors:** Laura A. Leuenberger, Joseph S. Bednash, Evangeline Schott, Carmen Mikacenic, Eric D. Morrell, Carson Richardson, Lynn A. Fussner, Jeffrey C. Horowitz, Rama K. Mallampalli, Saame Raza Shaikh, Robert M. Tighe, Kymberly M. Gowdy

**Author notes:** **Corresponding Author**: Laura A Leuenberger, 241 W 11^th^ Avenue, Suite 5082, Columbus, OH 43201.

## Abstract

Acute Respiratory Distress Syndrome (ARDS), a heterogeneous syndrome of hypoxic respiratory failure secondary to dysregulated pulmonary inflammation, is caused by diverse insults. Because of this heterogeneity, mechanisms and treatments are difficult to study. As a treatment, n-3 polyunsaturated fatty acid (PUFA) supplementation has had mixed results. PUFAs and downstream oxylipins are important to pulmonary inflammation but are not well defined in ARDS. We hypothesized that differences in fatty acid metabolism, as measured by levels of n-3 and n-6 PUFAs and oxylipins, are associated with differences in ARDS outcomes, ARDS causes, and inflammation.

To test this, PUFAs/oxylipins were measured by LC MS/MS in plasma samples from 90 patients with ARDS. Inflammatory cytokines (IL-6, IL-8) were measured by ELISA. Multivariate linear regressions modeled the relationship between PUFAs/oxylipins, inflammation, and ARDS mortality, severity and cause. Multiple n-3 and n-6 PUFA derived oxylipins were decreased in severe ARDS. We did not detect differences in PUFAs/oxylipins by mortality. PUFAs/oxylipins varied by cause of ARDS, especially between patients with sepsis and those with trauma. Furthermore, specific oxylipins were associated with IL-6 and IL-8. Based on our identification of oxylipins that vary by disease severity and injury, these metabolites should be considered as potential biomarkers and mechanisms of lung repair biomarkers. Furthermore, differences in PUFAs did not directly correlate with changes in oxylipins, suggesting differences in lipid metabolism by etiology of injury. Further consideration of differences in lipid metabolism in ARDS could identify potential subgroups that could benefit from n-3 PUFA supplementation or other therapies.

## Introduction

Acute Respiratory Distress Syndrome (ARDS) is a heterogeneous syndrome of severe lung injury caused by direct injury (i.e. bacterial or viral pneumonia, aspiration) or indirect injury (i.e. sepsis, massive transfusion, pancreatitis) (1). Heterogeneous injuries may cause distinct inflammatory mechanisms, complicating ARDS research and treatment. While hypo- and hyperinflammatory ARDS subphenotypes have been proposed to explain inflammation-driven outcomes (2), these subphenotypes do not consider etiology. Therefore, understanding how disparate causes of ARDS influence inflammation may help to identify molecular mechanisms and potential treatments.

Administration of n-3 (omega-3) polyunsaturated fatty acids (PUFAs) and their metabolites improves outcomes in animal models of sepsis and ARDS, yet clinical trials in humans show inconsistent results (3). For example, one randomized controlled trial supplementing n-3 PUFAs eicosapentaenoic acid (EPA) and docosahexaenoic acid (DHA) in sepsis- or trauma-related lung injury did not find differences in pulmonary or systemic inflammation (4). Conversely, a randomized controlled trial supplementing n-3 PUFAs EPA and DHA, n-6 PUFA gamma-linolenic acid (GLA), and antioxidants observed differences in inflammatory trajectories in sepsis-related ARDS (5). Adding to this evidence, PUFAs and their signaling metabolites, termed ‘oxylipins’ (i.e. oxygenated fatty acids), are known regulators of inflammation (6). Oxylipins generated from PUFAs can be synthesized nonenzymatically or enzymatically via lipoxygenases (LOX), cyclooxygenases (COX), and cytochromes (CYPs). PUFAs and oxylipins are altered in infectious and inflammatory lung diseases such as asthma (7), chronic obstructive pulmonary disease (COPD) (8, 9), COVID-19 (10, 11) and ARDS (12). However, the relationships between n-3 and n-6 PUFA metabolism, ARDS etiology, and ARDS-associated inflammation have not been determined.

We hypothesized that there are distinct patterns of n-3 and n-6 PUFAs and oxylipins that correlate with mortality, ARDS severity, ARDS etiology, and inflammation. To test this hypothesis, we measured PUFAs and oxylipins in 90 ARDS subjects and correlated with clinical outcomes. While PUFA and oxylipin levels did not predict mortality, select oxylipins were decreased in severe ARDS. Plasma PUFA and oxylipins significantly changed based on ARDS etiology. Additionally, PUFAs and oxylipins correlated with inflammatory cytokines interleukin (IL)-6 and IL-8. Collectively, our findings demonstrate a unique and previously undescribed role of PUFAs/oxylipins in ARDS.

## Materials and Methods

### Study design and patient population

Patient samples from two separate biorepositories were used for this study. The first biorepository was the Ohio State University Intensive Care Unit Registry (IRB 2020H0175: The Ohio State University Intensive Care Unit Registry (BuckICU), 2023H0138: Critical Illness Phenotyping of Existing Biospecimens) an ongoing prospective single-center cohort of adult patients hospitalized with ARDS or critical illness started in 2020. Patients were screened on admission to the intensive care unit, and consent was obtained within 48 hours of admission. Enrollment criteria included either (a) respiratory failure, defined as a need for supplemental respiratory support above baseline, including nasal cannula, heated high flow nasal cannula, non-invasive positive pressure ventilation, or invasive mechanical ventilation to maintain an oxygen saturation greater than 92%, or (b) met sepsis criteria by Sepsis-3 guidelines (13). The second biorepository was the University of Washington (IRB #3412: University of Washington Human Subjects Division). Samples were collected from a prospectively enrolled cohort of critically ill adults with acute hypoxemic respiratory failure on mechanical ventilation between September 2020 and December 2023. Enrollment occurred within 96 hours of ICU admission. All study protocols were approved and monitored by the respective local institutional review boards (IRB) at The Ohio State University and University of Washington and abided by the Declaration of Helsinki principles.

In both cohorts, ARDS diagnosis was adjudicated by two pulmonary/critical care physicians, using the new global definition of ARDS (14). At that time of adjudication, a clinical decision of the cause of the ARDS was assigned. The etiologies of ARDS were defined as sepsis, pneumonia, aspiration, trauma, and other. In both registries, recognizing that ARDS is frequently multifactorial, a case could be assigned up to two causes of ARDS. Therefore, ARDS etiology groups overlap in this analysis. For this study, a total of 90 patients (all intubated and mechanically ventilated) from these two registries met ARDS criteria and had available samples (59 from Ohio State and 31 from University of Washington). Given the prior collection of these samples, blinding was not possible in the selection of samples, however, all laboratory analyses were completed in batches without the knowledge of any clinical data and thus run in a blinded fashion. The sex and gender of both biorepositories reflected the general heterogeneous ICU population, and samples used for the analysis replicated those populations. For both cohorts, patient data was stored securely in REDCap.

### Blood collection and processing

Peripheral blood samples were collected in citrate vacutainer tubes (BD Biosciences, Franklin Lake, New Jersey), centrifuged at 1200 x g for 10 min and then stored at -80°C.

### Fatty acid and oxylipin measurements

Patient plasma samples stored at -80°C from both repositories that had not undergone a prior freeze/thaw cycle were processed with 1:1 ratio of 75% ethanol for lipid extraction. Samples were again stored at -80 °C until the time of processing. LC-MS/MS was performed by the Lipidomics Core Facility at Wayne State University as previously published (15). Briefly, 200 μl of sample were spiked with 5 ng of the internal standards and then diluted to 1 ml with 15% methanol in water and purified on C18 solid-phase extraction cartridges. The diluted sample (with standards) was applied to the cartridge with 2mL of 15% methanol and 2 mL of hexane and dried in a vacuum for 30s. The cartridge was eluted with 0.5 ml methanol with 0.1% formic acid into LC-MS autosample vials. The elute was dried under nitrogen then reconstituted with 25 μl of methanol. For analysis, the sample was brought to room temperature, 25 μl of 25 mM aqueous ammonium acetate was added, mixed and loaded into the autosampler at 15°C. HPLC was performed on a C18 column, as previously published (15). Plasma concentration of each detected analyte was calculated by dividing the detected quantities (in nanograms) with their corresponding molecular masses and reported as nanomolar. This lipidomic analysis quantified n-3 and n-6 PUFAs (i.e. EPA, DHA, Docosapentaenoic Acid (DPA), Arachidonic Acid (AA), and Linoleic Acid (LA)) and targeted oxylipins (>120 fatty acid derived metabolites). Due to technical limitations, n-3 alpha-linolenic acid (ALA) and n-6 gamma-linolenic acid (GLA) measurements were combined. A list of measured lipids is shown in Supplemental Table 1.

### Measurements of systemic inflammation

Plasma samples were used to measure IL-6 and IL-8 using the V-PLEX Human Proinflammatory Panel 1 Kit (Catalog number K15049D, Meso Scale Discovery, Rockville, MD).

### Statistical Analysis

To correlate clinical outcomes, a series of both univariable and multivariable linear regression models were used to define associations of individual metabolites with mortality and disease severity (primary outcomes). Mortality was defined by in-hospital mortality. Disease severity was defined using the Berlin Criteria PaO_2_/FiO_2_ ratio of mild (200-300), moderate (100-200) and severe (<100) (16). Given the paucity of participants who met “mild” criteria (N=6), severity and associated responses were analyzed as “Mild/Moderate” versus “Severe.” Missing data within the metabolites were imputed using the K-nearest neighbor truncation algorithm (17) and any metabolites with >25% missing data were excluded from analysis. The metabolite data were log^2^-transformed to handle the skewed distributions across the sample. The Benjamini-Hochberg false discovery rate was implemented to account for multiple comparisons with a significance threshold of q < 0.1 (18).

Multivariate linear regression was then used to describe the relationships of PUFAs/oxylipins to the secondary outcome variables of “ARDS cause” (categories defined above), comparing all causes to sepsis related ARDS, as this is best described etiology in the both the PUFA/oxylipin and ARDS literature. Inflammation, measured by cytokines IL-6 and IL-8, was used as a secondary outcome variable. The relationships between IL-6/IL-8 and PUFAs/oxylipins were evaluated via separate linear regression models using the same imputation algorithm and log^2^-transformed values. Given known alterations of lipid metabolism by age (19), sex (20), statin use (21), diabetes (22), and cancer (23), the model was adjusted for these variables, as well as for clinical site, race, and COVID-19 status. Adjusted and unadjusted models had similar results and thus only the unadjusted models are shown. Analysis was performed using R version 4.5.1 and GraphPad Prism 10.

## Results

### Baseline Cohort Characteristics

Table 1 shows the baseline characteristics of this cohort, which had a median age of 55 years, and was predominantly male (60%, N=54), White (77%, N=69) and non-Hispanic/Latino (95%, N=81). ARDS etiologies included pneumonia (67.7%), sepsis (61.1%), aspiration (24.4%), trauma (13.3%), and “other” causes (13.3%) such as transfusion-related or inhalational injuries. As our cohort enrolled during COVID-19 pandemic, 37% (N=33) of subjects tested positive for SARS-CoV-2. Patients who died during the index hospitalization were older (*P*=0.035), and more commonly COVID-19 positive (*P*=0.029).

**Table 1:** Baseline Characteristics of Study Population of Patients with Acute Respiratory Distress Syndrome (ARDS) by Hospital Mortality. Frequencies are presented as number (%) unless otherwise indicated. Continuous variables are presented as Median (Q1, Q3). BMI: Body Mass Index (kg/m^2^). Race and ethnicity were identified from the patient chart, as reported by patients or surrogates. COPD: Chronic Obstructive Pulmonary Disease. ILD: Interstitial Lung Disease. COVID-19: Coronavirus Disease 2019. *P*-values indicate difference between Survived and Died population, as measured by Mann-Whitney U test (continuous variables) or Fisher’s Exact test (categorical variables), with significance of *P*<0.05.

| Characteristic |  | Overall<br>N = 90 | Died<br>N = 38 | Survived<br>N = 52 | P-Value |
| --- | --- | --- | --- | --- | --- |
| Age (Median (Q1, Q3)) |  | 55 (41, 66) | 60 (48, 70) | 53 (33, 65) | 0.035 |
| Female Sex |  | 36 (40%) | 15 (39%) | 21 (40%) | >0.999 |
| BMI (Median (Q1, Q3)) |  | 29 (25, 36) | 29 (25, 34) | 30 (25, 36) | 0.722 |
| Race | White | 69 (77%) | 28 (74%) | 41 (79%) | 0.619 |
|  | Black or African American | 15 (17%) | 8 (21%) | 7 (13%) | 0.397 |
|  | Other | 6 (6.7%) | 2 (5.3%) | 4 (7.7%) | >.0.999 |
| Ethnicity | Not Hispanic or Latino | 81 (98%) | 34 (97%) | 47 (98%) | >.0.999 |
|  | Hispanic or Latino | 2 (2.4%) | 1 (2.9%) | 1 (2.1%) | >.0.999 |
|  | Unknown | 7 (7.7%) | 3 (7.9%) | 4 (7.7%) | >.0.999 |
| Past medical history | History of COPD | 16 (18%) | 7 (18%) | 9 (17%) | >.0.999 |
|  | History of ILD | 2 (2.2%) | 0 (0%) | 2 (3.8%) | 0.507 |
|  | History of Asthma | 15 (17%) | 6 (16%) | 9 (17%) | >.0.999 |
|  | Home O2 | 7 (7.8%) | 4 (11%) | 3 (5.8%) | 0.450 |
|  | History of Diabetes | 26 (29%) | 13 (34%) | 13 (25%) | 0.357 |
|  | Active Cancer | 15 (17%) | 8 (21%) | 7 (13%) | 0.397 |
|  | Prior Statin Use | 17 (19%) | 9 (24%) | 8 (15%) | 0.415 |
| COVID-19 positive |  | 33 (37%) | 19 (50%) | 14 (27%) | 0.029 |
| ARDS Severity | Mild | 6 (6.7%) | 1 (2.6%) | 5 (9.6%) | 0.395 |
|  | Moderate | 40 (44%) | 14 (37%) | 26 (50%) | 0.284 |
|  | Severe | 44 (49%) | 23 (61%) | 21 (40%) | 0.087 |
| Causes of ARDS | Pneumonia | 61 (67.7%) | 28 (73.7%) | 33 (63.4%) | 0.365 |
|  | Sepsis | 55 (61.1%) | 26 (68.4%) | 29 (55.8%) | 0.276 |
|  | Aspiration | 22 (24.4%) | 10 (26.3%) | 12 (23.1%) | 0.724 |
|  | Trauma | 12 (13.3%) | 2 (5.3%) | 10 (19.2%) | 0.065 |
|  | Other | 12 (13.3%) | 5 (13.2%) | 7 (13.4%) | 0.967 |

### Peripheral levels of n-3 and n-6 PUFAs and oxylipins do not predict ARDS mortality

We first determined if specific n-3 or n-6 PUFAs or downstream oxylipins were associated with ARDS mortality. When comparing plasma PUFAs and oxylipins to mortality, there were no statistical differences in any n-3 or n-6 PUFAs (Figure 1), or their downstream oxylipins (Supplemental Table 2) when stratified by in-hospital mortality.

**Figure 1:**
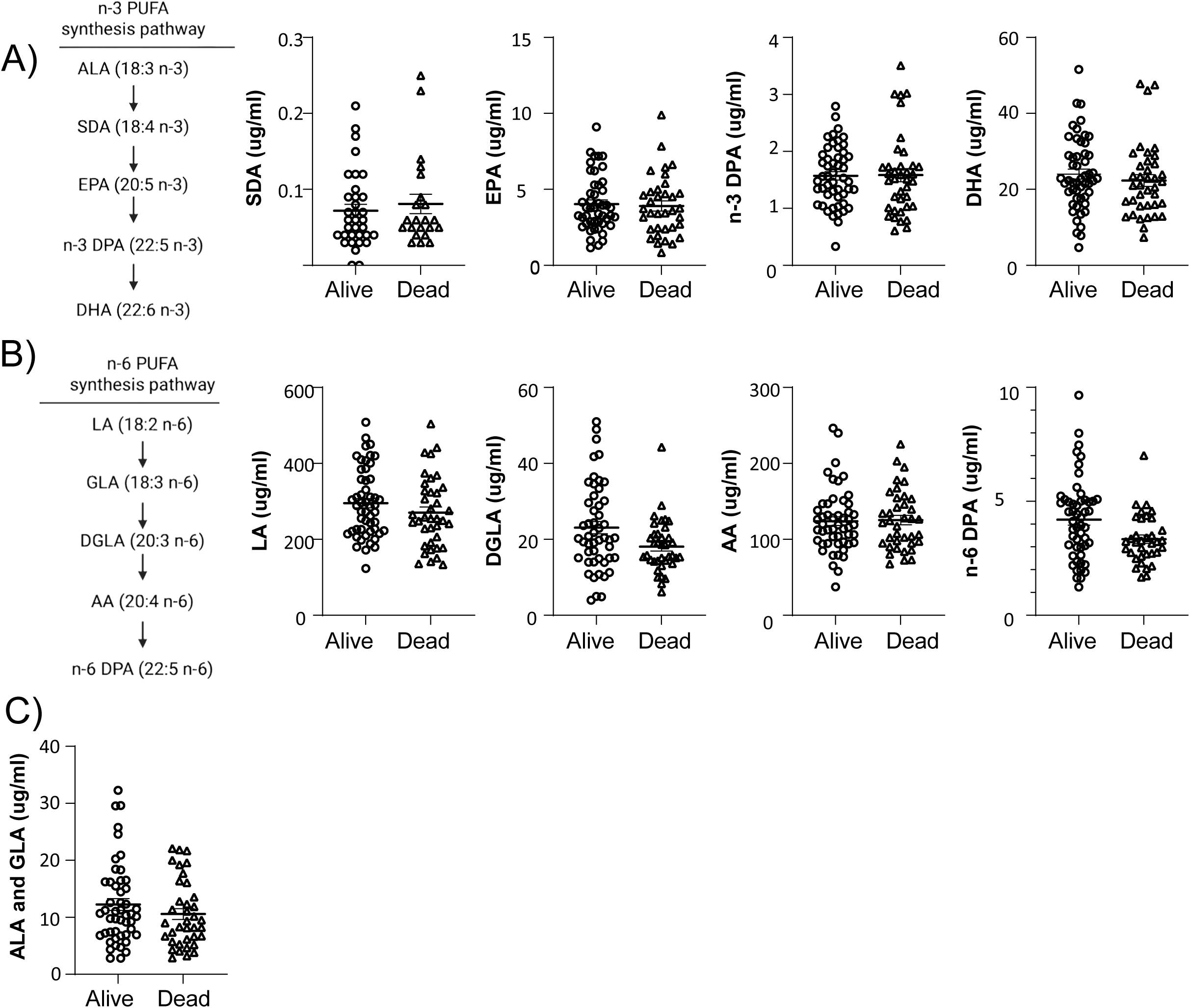
Circulating levels of n-3 and n-6 polyunsaturated fatty acids (PUFAs) are not associated with in-hospital mortality in ARDS patients. Plasma was collected from patients within 96 h of hospital admission and total fatty acid levels were measured by LC-MS/MS. Fatty acid levels were separated by survived versus died during the hospital admission. (**A**) Diagram of n-3 PUFA synthesis pathway and levels of n-3 PUFAs in plasma including stearidonic acid (SDA), eicosapentaenoic acid (EPA), n-3 docosapentaenoic acid (DPA), and docosahexaenoic acid (DHA). (**B**) Diagram of n-6 PUFA synthesis pathway and n-6 PUFA levels in plasma including linoleic acid (LA), dihomo-gamma-linolenic acid (DGLA), arachidonic acid (AA), and n-6 DPA. (**C**) Plasma levels of combined alpha (n-3) and gamma (n-6) linoleic acid (ALA and GLA). N = 38 Died, 52 Alive.

### Select n-3 and n-6 PUFA derived oxylipins in plasma predict ARDS severity

We next investigated if levels of circulating PUFAs/oxylipins were associated with ARDS severity using multivariate linear regression. Multiple oxylipins were decreased in severe ARDS, specifically n-3 PUFA/EPA derived Resolvin E1 (RvE1) (Figure 2A) and prostaglandin (PG) 15d-D12,14-PGJ3 (Figure 2B) as well as n-6 PUFA /AA derived 15d-D12,14-PGJ2 (Figure 2C), and n-6 PUFA/linoleic acid (LA) oxylipins 9,10-DiHOME (Figure 2D), and 12,13-DiHOME (Figure 2E). However, similar to mortality, there were no differences in the peripheral levels of n-3 or n-6 PUFAs by disease severity (data not shown).

**Figure 2:**
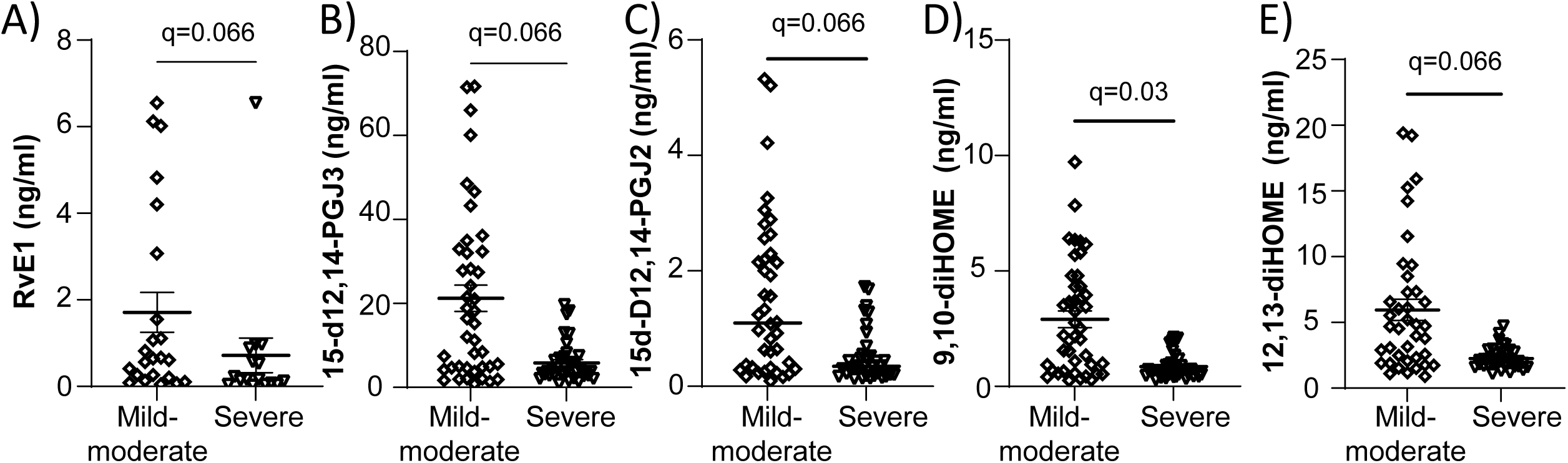
Oxylipins are decreased in the plasma of patients with severe ARDS. Plasma was collected from patients within 96 h of hospital admission and oxylipin levels were measured by LC-MS/MS. Oxylipins were separated by those with Mild-Moderate ARDS versus those with Severe ARDS. Categories were determined by Berlin Criteria, with mild-moderate PaO_2_:FiO_2_ =100-300 and severe with PaO_2_:FiO_2_ <100. Plasma concentrations of EPA-derived oxylipins (**A**) Resolvin E1 (RvE1) and (**B**) 15-d12,14-PGJ3, Arachidonic Acid derived (**C**) 15d-D12,14-PGJ2 and Linoleic Acid derived (**D**) 9,10-diHOME and (**E**) 12,13-diHOME. Graphed are the absolute concentrations without any imputed values for visual display. Significance identified by multivariable regression models comparing individual metabolites and disease severity, with Benjamini-Hochberg false discovery rate accounting for multiple comparisons with a significance threshold of *Q*<0.1. N= 46 mild-moderate, 44 severe.

### Peripheral PUFAs and oxylipins vary by ARDS etiology

Given the relationship of RvE1 to disease severity and preclinical data regarding its impact on lung injury (24), we first investigated alterations in EPA and EPA metabolites along this pathway by ARDS cause (Figure 3A). Plasma EPA was decreased in sepsis-related ARDS when compared to trauma associated ARDS (Figure 3B). Other differences included increased EPA derived oxylipins 12-hydroxyeicosapentaenoic acid (HEPE) in the sepsis group (*P*<0.05) and 15-HEPE in both sepsis (*P*<0.001) and pneumonia (*P*<0.001) when compared to trauma (Figure 3D-E). There were no differences in RvE1 levels by ARDS etiology (Figure 3G).

**Figure 3:**
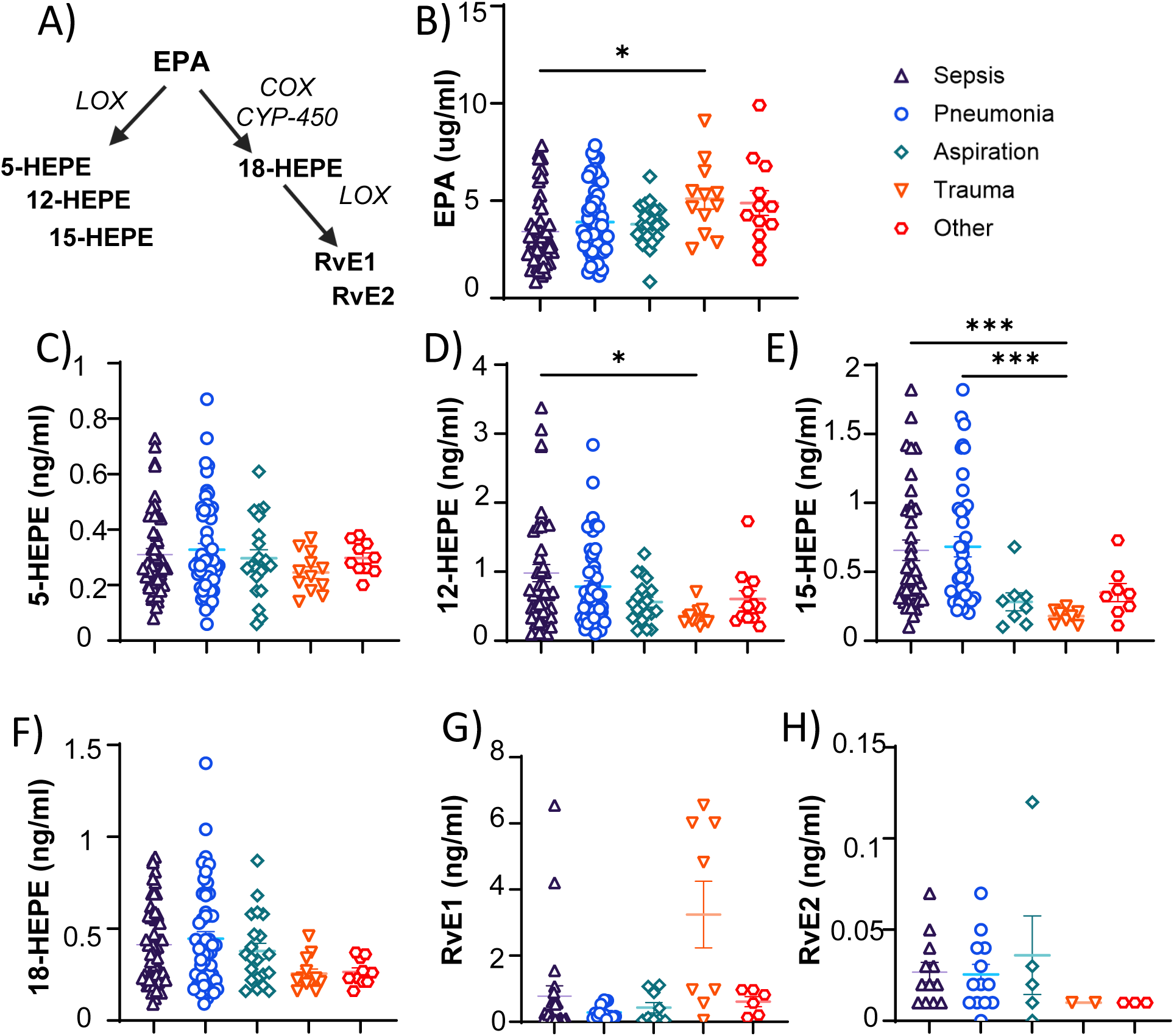
Eicosapentaenoic Acid (EPA) and EPA-derived mediators differ by ARDS etiology. Plasma was collected from patients within 96 h of hospital admission and total fatty acid and oxylipin levels were measured by LC-MS/MS. Cause of ARDS was determined by the research investigators and could include up to two causes. (**A**) Simplified diagram of a subset of Eicosapentaenoic Acid metabolism, including relevant enzymes lipoxygenases (LOX), cyclooxygenases (COX), and cytochrome P450 (CYP-450). Plasma concentrations of (**B**) EPA, (**C**) 5-Hydroxyeicosapentaenoic acid (HEPE), (**D**) 12-HEPE, (**E**) 15-HEPE, and (**F**) 18-HEPE, (**G**) Resolvin E1 (RvE1), (**H**) RvE2, compared across disease groups. Outliers removed by ROUT (Q=1%), Analysis by one-way ANOVA (Kruskal-Wallis), N=12-61/group. \**P*<0.05, \*\*\**P*<0.001

Next, we analyzed PUFAs and oxylipins by ARDS cause using multivariable regression modeling (significance = *Q*-value <0.1) and made all comparisons to ARDS secondary to sepsis. Regarding PUFAs, ALA/GLA were increased in both aspiration (*Q*=0.009) and trauma (*Q*<0.001), LA was increased in aspiration (*Q*=0.015), trauma (*Q*<0.001) and other (*Q*<0.001), dihomo-gamma-linolenic acid (DGLA) was increased in trauma (*Q*=0.023), and n-6 docosapentaenoic acid (DPA) was increased in other (*Q*=0.06) (Figure 4A, Supplemental Table 3). However, unlike in our initial analysis (Figure 3), differences in EPA were not statistically significant in this model.

**Figure 4:**
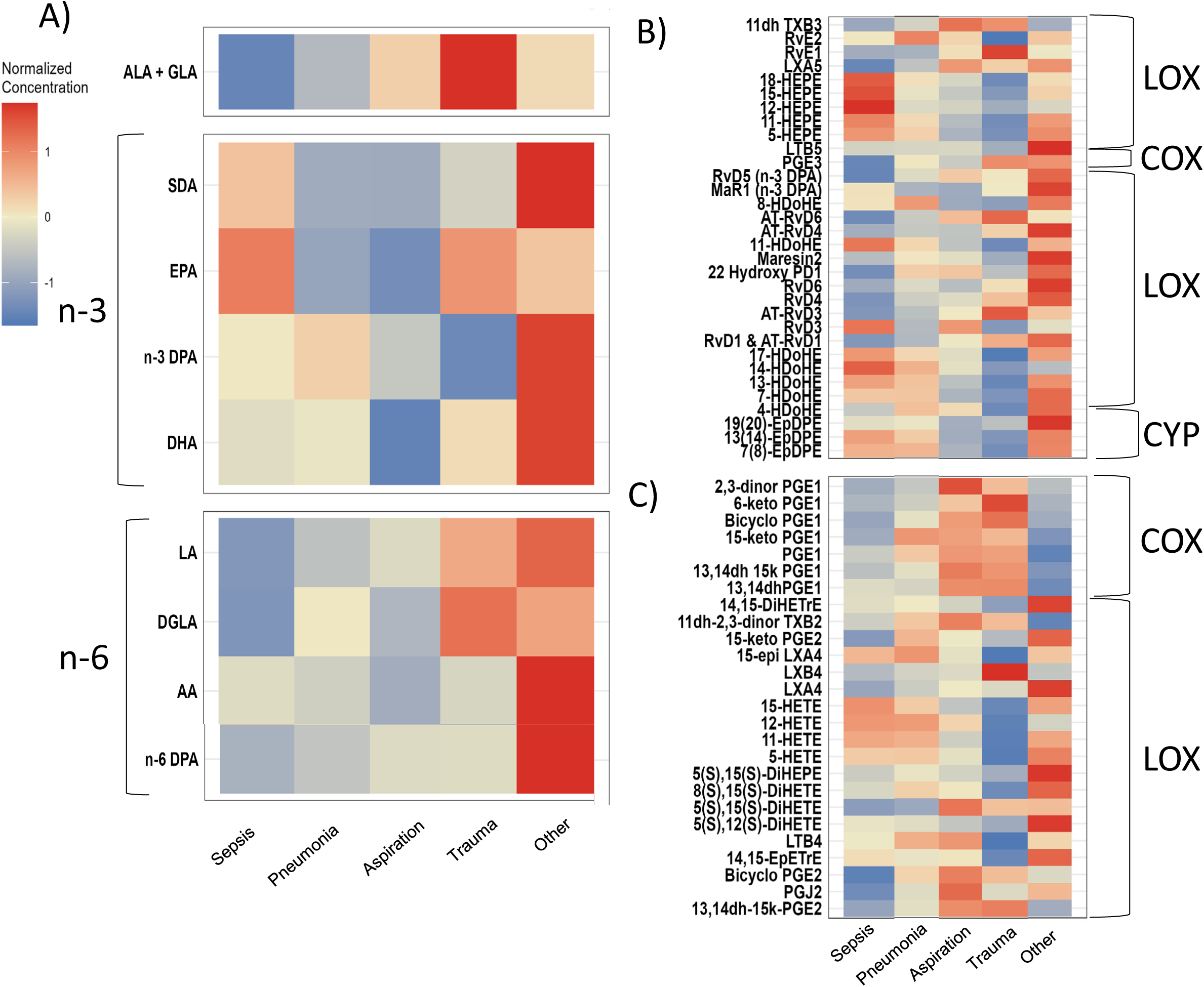
Plasma polyunsaturated fatty acids (PUFAs) and oxylipins levels are altered in different ARDS etiologies. Plasma was collected from patients within 96 h of hospital admission and fatty acid and oxylipin levels were measured by LC-MS/MS. Metabolites were analyzed based on ARDS etiology. Etiology of ARDS was determined retrospectively based on chart review by original study investigators; each patient could have up to two causes of ARDS that were included in this analysis. PUFAs and oxylipins were transformed by log^2^-fold change to account for outliers and then normalized to allow comparison between metabolites; color represents relative concentration (unitless given normalization). Indicated are the parent PUFA (n-3 or n-6), and oxylipin enzymatic pathways: lipoxygenase (LOX), cyclooxygenase (COX) and cytochrome P450 (CYP). Heat maps demonstrate (**A**) fatty acids normalized concentrations of alpha-linoleic acid (ALA), gamma-linoleic Acid (GLA), stearidonic acid (SDA), eicosapentaenoic acid (EPA), n-3 docosapentaenoic acid (DPA), and docosahexaenoic acid (DHA), linoleic acid (LA), dihomo-gamma-linolenic acid (DGLA), arachidonic acid (AA), and n-6 DPA. (**B**) Heat Map of n-3 derived oxylipins and (**C**) Heat Map of n-6 derived oxylipins. Abbreviations defined in e-Table 1. Plasma samples from 90 patients with ARDS were analyzed, N=12-61/group.

Generally thought to be proinflammatory, multiple PGs were increased in all groups compared to patients with sepsis. Increased AA-derived PGs included 8-isoPGF2a (Aspiration *Q*<0.001, Other *Q*=0.013, Pneumonia *Q*=0.025, Trauma *Q*<0.001), bicyclo-PGE2 (Aspiration *Q*=<0.001, Pneumonia *Q*=0.018, Trauma Q=0.024), 13,14-dh-15k-PGE2 (Aspiration *Q*=0.006, Trauma *Q*=0.017), 15-keto-PGF2a (Trauma *Q*=0.095), d12-PGJ2 (Aspiration *Q*=0.003, Pneumonia *Q*=0.043, Trauma *Q*=0.018), PGJ2 (Aspiration *Q*<0.001, Other *Q*=0.067, Trauma *Q*=0.026), 15d12,14-PGJ2 (Aspiration *Q*<0.001, Other *Q*=0.067, Trauma *Q*=0.026), 6-keto-PGE1 (Aspiration *Q*=0.058, Trauma *Q*=0.011), 6,15-diketo-PGFa (Aspiration *Q*<0.001, Trauma *Q*=0.006) and 19(R)-OH-PGF2a (Other *Q*=0.024, Trauma *Q*=0.017) (Figure 4C, Supplemental Table 4).

DGLA-specific PG increased include 2,3,-dinor PGE1 (Aspiration, *Q*<0.001, Trauma *Q*=0.028), 15-ketoPGE1 (Aspiration *Q*=0.053, Pneumonia *Q*=0.053) and PGF1a (Aspiration *Q*<0.001, Pneumonia *Q*=0.07, Trauma *Q*<0.001). Specific to aspiration were increases in DLGA-derived 13,14-dh-15k-PGE1 (*Q*=0.086), PGJ2 (*Q*=0.045). EPA derived PGs were also increased, including 15d,d12,14-PGJ3 (Aspiration *Q*<0.001, Other *Q*=0.07, Trauma *Q*<0.001).

Thromboxanes (TX) were also elevated compared to sepsis, including AA-derived 2,3,dinor-TX*B2* (Trauma *Q*=0.007) and 11,dh-TXB3 (Aspiration *Q*<0.001, Trauma *Q*=0.007). Patients with aspiration had increases in 11,dh-2,3,-dinor-TXB2 (*Q*=0.082). However, other AA-derived metabolites were decreased. In the trauma cohort, decreased AA-derived hydroxyeicosatetraenoic acids (HETE) included *8(S),15(S)-*diHETE (*Q*=0.067), 5-HETE (*Q*=0.007), 8-HETE (*Q*=0.007), 11-HETE (*Q*=0.002),

12- HETE (*Q*=0.023), 15-HETE (*Q*=0.019), and 20-HETE (*Q*=0.017). Both the aspiration and trauma groups had decreased 5(S),15(S)-diHETE (Aspiration *Q*<0.001, Trauma *Q*=0.036), and trauma specifically had decreased 5,6-di-HETrE (*Q*=0.025), another AA derived metabolite. Also decreased was 14,15-EPETrE (Aspiration *Q*=0.006, Pneumonia *Q*=0.082, Trauma *Q*<0.001). The other group differed from sepsis with increased AA-derived lipoxin (LX) LXA4 (*Q*=0.085) (Figure 4C).

Compared to sepsis, EPA-derived metabolites were decreased in trauma patients including 11-HEPE (*Q*=0.07) and 18*-*HEPE (*Q*=0.036), RvE2 (*Q*=0.045), while 15*-*HEPE (Aspiration *Q*=0.011, Pneumonia *Q*=0.072, Trauma *Q*<0.001), 12*-*HEPE (Aspiration *Q*=0.086, Trauma *Q*=0.029) and 17,18-EPETE (Aspiration *Q*=0.054, Trauma *Q*=0.003) were decreased in multiple cohorts. In contrast, other EPA derived metabolites were increased including RvE1 (Aspiration *Q*=0.038, Trauma *Q*=0.003) and LXA5 in aspiration (*Q*=0.013). The other category also had increases in EPA derived leukotriene (LT) LTB5 (*Q*=0.082) and HEPE 5(S),15(S)-diHEPE (*Q*=0.024) (Figure 4B, Supplemental Table 4).

DHA metabolites, Resolvin Ds (RvDs) were increased in multiple causes compared to sepsis, including 8-oxo-RVD1 (Aspiration *Q*=0.89), RvD1/AT-RvD1 (Aspiration *Q*=0.07, Other *Q*=0.033, Trauma *Q*=0.029), RvD4 (Other *Q*=0.036, Trauma *Q*=0.064), RvD6 (Aspiration *Q*=0.031, Trauma *Q*=0.017). DHA derived Protectin (PD) 22-OH-PD1 was also increased (Aspiration *Q*=0.041, Other *Q*=0.067, Pneumonia *Q*=0.063). N-3 DPA derived RvD5 was also increased in multiple groups compared to sepsis (Aspiration *Q*=0.019, Other *Q*=0.028, Trauma *Q*=0.042, *Q*=0.095). DHA specific mediators decreased in trauma patients included hydroxydocosahexaenoic acid (HDoHEs) 11-HDoHE (*Q*=0.017),

13- HDoHE (*Q*=0.045), 14-HDoHE (*Q*=0.017), 16-HDoHE (*Q*=0.07), 17-HDoHE (*Q*=0.005) Decreases in 19,20-diHDoPE (*Q*=0.029), and 7,8-EPDPE (Aspiration *Q*=0.082, Trauma *Q*=0.024). In contrast, the other group had increases in DHA metabolites 20-HDoHE (*Q*=0.058) and Maresin2 (*Q*=0.045) (Figure 4B, Supplemental Table 4).

LA-derived diHOMEs were increased compared to sepsis including 9,10-diHOME (Aspiration *Q*=0.001, Other *Q*=0.025, Trauma *Q*=0.002) and 12,13-diHOME (Aspiration *Q*=0.006, Other *Q*=0.025). Aspiration specific differences included increased LA derived hydroxyoctadecatrienoic acids (HOTrE) including 9-HOTrE (*Q*=0.007) and 13*-*HOTrE (*Q*=0.006) and increased AA derived hydroxyoctadecadienoic acid (HODE) 9-HODE (*Q*=0.045), 13-HODE (*Q*=0.07), and AA derived 15-oxo-ETE (Aspiration *Q*=0.001, Pneumonia *Q*=0.07). All significant differences are detailed in Supplemental Table 6.

### Specific PUFAs and oxylipins in the plasma of patients with ARDS correlate with inflammation

To consider these findings within the context of hyper- and hypo-inflammatory subphenotypes of ARDS (2), we compared the relationship of plasma IL-6 and IL-8 levels to PUFAs and oxylipins. Plasma levels of AA negatively correlated with IL-6 (*Q*=0.014) (Supplemental Table 5). All other n-3 and n-6 PUFAs had a negative trend with IL-6 but did not reach significance (Figure 5). When compared to plasma IL-8 levels, circulating PUFAs n-3 DPA (*Q*=0.03) and n-6 PUFAs AA (*Q*=0.013) and DGLA (*Q*=0.085) were negatively correlated (Supplemental Table 5).

**Figure 5:**
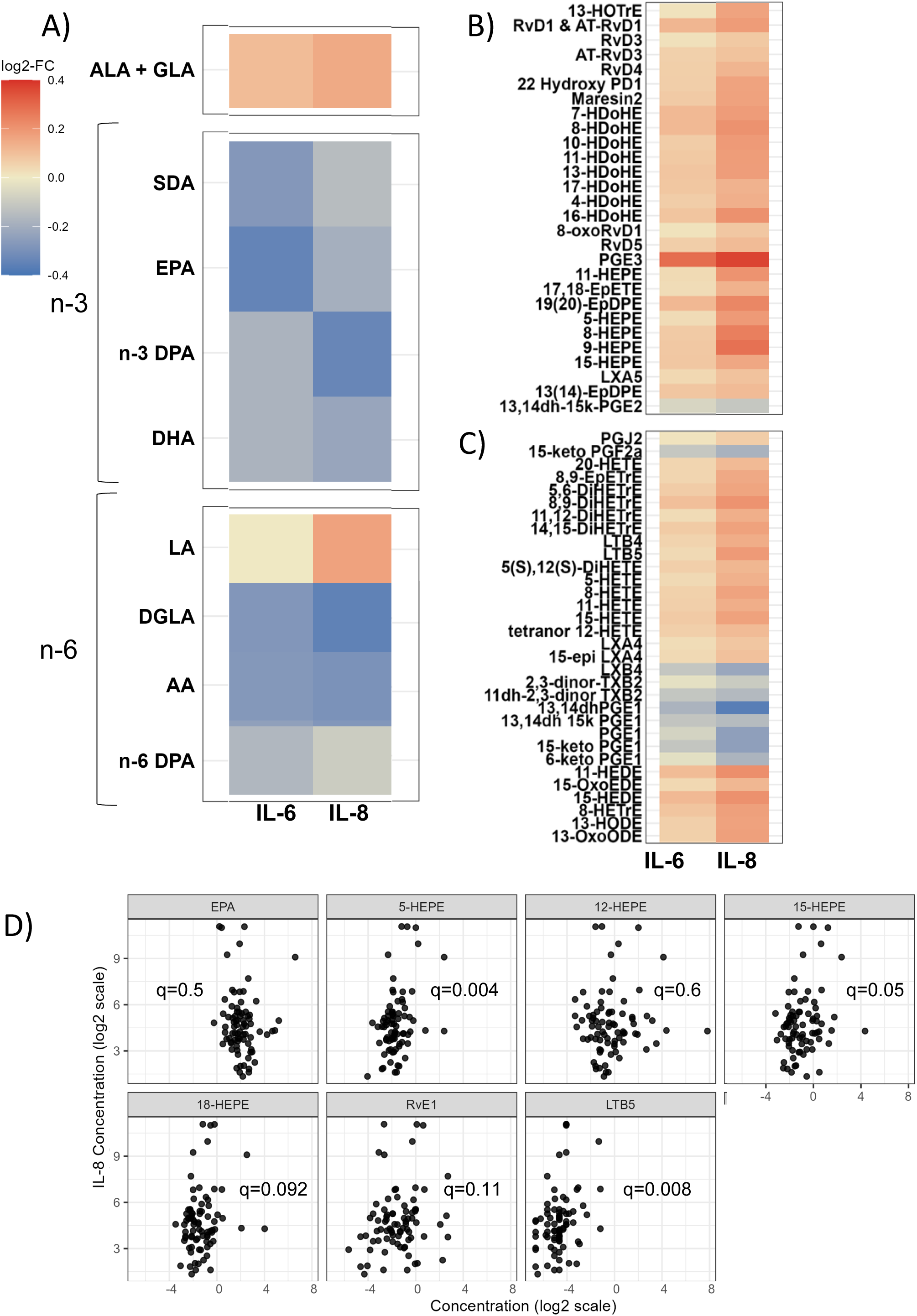
Plasma PUFA and oxylipins are associated with circulating proinflammatory cytokines. Plasma was collected from patients within 96 h of hospital admission and total fatty acid and oxylipin levels were measured by LC-MS/MS. IL-6 and IL-8 were measured by multiplex ELISA. Colors represent normalized concentration of (**A**) fatty acids alpha-linolenic Acid (ALA), gamma-linolenic Acid (GLA), stearidonic acid (SDA), eicosapentaenoic acid (EPA), n-3 docosapentaenoic acid (DPA), docosahexaenoic acid (DHA), linoleic acid (LA), dihomo-gamma-linolenic acid (DGLA), arachidonic acid (AA), and n-6 DPA. (**B**) n-3 derived oxylipins and (**C**) n-6 derived oxylipins compared to cytokine levels, normalized by log^2^-fold change to account for outliers. (**D**) Scatter plot of EPA and EPA-derived oxylipins plotting log2 transformed IL-8 concentrations (x-axis) and log^2^ metabolite concentrations (y-axis). The Benjamini-Hochberg false discovery rate accounted for multiple comparisons with a significance threshold of q< 0.1. N=80.

When evaluating the correlation between inflammatory cytokines and oxylipins, both n-3 and n-6 PUFA-derived oxylipins were generally positively correlated with IL-6 and IL-8, suggesting increased response to inflammation (Figure 5B-5C). There were also, however, n-6 PUFA derived oxylipins, primarily PG and TXs, negatively correlated with inflammation (Figures 5C). IL-6 was negatively associated with oxylipins including 13,14-dh-PGE, 13,14-dh-15k-PGE, PGE1, 15-ketoPGE1, 13,14-dh-15k-PGE2, and 15-keto-PGF2, as well as 11-dh,2,3,dinor-TXB2 (Supplemental Table 6). Oxylipins positively correlated with IL-6 included PGE3, 19,20 EpDPE, and 8,9-di-HETrE. IL-6 was positively correlated with DHA derived 7-HDoHe, 20-HDoHE, AT-RvD1 and AT-RvD4. IL-8 plasma levels were negatively associated with prostaglandins 13,14-dh-PGE, PGE1, 15-keto-PGE1, 13,14-dh-15k-PGE, 15-keto-PGF2a, 11,dh-2,3-dinor-TXB2, LXB4, 6-keto-PGE1, 13,14-dh-15k-PGE2,2,3-dinor-TXB2. PGE3 was positively correlated with IL-8, as were PGE2, PGJ2, LTB4 and LTB5. AA derived HETEs (5-, 8-, 11, 15, and 20), EPA derived HEPES (5-, 8-, 9-, 11-, 15-, 18-) (Figure 5D), and DHA derived HDoHEs (4-, 7-, 8-, 10, 11-, 13-, 16-, 17-) were all positively correlated with IL-8. Lastly, DHA-derived resolvins (AT-RvD1, RvD3, AT-RvD3, 8-oxoRvD1 RVD4, AT-RvD4, and RVD6) were all positively correlated with IL-8 (Supplemental Table 6).

## Discussion

To better understand ARDS heterogeneity, we characterized PUFAs and their oxylipins in patients with ARDS from diverse causes. This study, the largest of its kind, identified select circulating n-3 and n-6 PUFAs and downstream oxylipins associated with ARDS severity and inflammation. Furthermore, we demonstrated that PUFA and oxylipin profiles are associated with etiologies of ARDS, suggesting differential roles for PUFA and oxylipins based on ARDS cause. We also found relationships between inflammation and PUFAs/oxylipins. Together, these data define PUFA metabolism as differentially regulated by various ARDS etiologies, which may predict inflammation and/or disease severity.

Our data show that plasma levels of EPA, LA, and AA-derived oxylipins were elevated in less severe ARDS, although circulating PUFAs and oxylipins did not predict mortality. Prior studies have shown decreases in both n-3 and n-6 PUFAs with ARDS (25), but little is known about the associations between PUFAs and ARDS outcomes. There are several potential mechanisms by which oxylipin levels may influence ARDS severity. EPA-derived RvE1, which we found to be decreased in severe ARDS, is known to improve alveolar fluid clearance (26) and to decrease inflammatory cytokines in animal models of pneumonia and aspiration (24). We speculate that RvE1 may play a similar role in mitigating ARDS severity. The LA-derived oxylipins that were decreased in severe disease, 9,10-diHOME and 12,13-diHOME, are well described as cardioprotective through CaMKII inhibition and decreased endoplasmic reticular stress (27). These LA-derived oxylipins have not been explored in the context of ARDS, but similar mechanisms may confer protective effects. Finally, we identified decreased levels of the oxylipin 5,D12-14-PGJ2 in patients with more severe ARDS, which is associated with asthma resolution (28) through its role activating the PPAR-gamma pathway (29). Together, our findings highlight the plausibility that specific PUFAs/oxylipins associated with ARDS severity may have a mechanistic role in ARDS pathobiology.

Next, we identified differences in specific oxylipins between patients with sepsis and those with multiple other etiologies of ARDS. Our sepsis findings are consistent with established literature, where both 12-HEPE and 15-HEPE are known to be increased in sepsis models (30). The differences in oxylipins by etiology could be related to the time from injury to ARDS, as oxylipin levels are known to be dynamic throughout inflammation and resolution (31). Our differences were most robust between patients with sepsis and those with trauma. While multiple prospective ARDS trials have targeted lipid pathways in sepsis-related ARDS cohorts (32, 33), our findings illustrate that the results may not be generalizable to other causes of ARDS. Overall, this is the first effort, to our knowledge, to identify phenotypes of ARDS by PUFAs/oxylipins, and adds to our understanding that differences in ARDS etiologies may drive differential inflammation.

A key novel finding of this study is the absence of an association between differences in PUFAs and their downstream oxylipins, perhaps identifying a cause for prior failed n-3 supplementation trials. These supplementation studies have been built on the idea that increased n-3 supplementation would increase downstream oxylipins, yet we did not see that reflected in these data. For example, while EPA-derived oxylipins were different based on etiology, EPA itself was not different between groups in our multivariate model. While increasing dietary n-3 PUFAs increases n-3 PUFA-derived oxylipins in mice (34) and healthy human subjects (35), our study suggests that PUFAs may not be metabolized the same in all injury types. We hypothesize this may be due to differential regulation of enzymatic pathways (LOX, COX, CYP450) by ARDS etiology driving differences in downstream oxylipin concentrations. Supporting this, a prior study of COVID-19 identified differences in oxylipins between mild and severe disease with increases in downstream metabolites specific to LOX-derived enzymes (11). Further clinical studies of PUFA supplementation may consider supplementing downstream oxylipins directly or targeting the mechanisms of PUFA metabolism.

Finally, we demonstrated that n-3 PUFA derived oxylipins were associated with pro-inflammatory cytokines IL-6 and IL-8. While our ability to directly compare the previously described hypo- and hyper-inflammatory sub-phenotypes was limited by clinical data and sample availability, using IL-6 and IL-8 as surrogates for hyperinflammatory subphenotypes suggests that there are associations between these markers of inflammation and oxylipin profiles. However, contrary to published subphenotypes, in this cohort, we did not see differences in mortality by IL-6 or IL-8 levels (data not shown), perhaps due to our smaller sample size. Even so, the relationship between these inflammatory markers and the n-3 derived oxylipins suggest a role for oxylipins as potential biomarkers in these models. Further investigation is needed to understand if specific oxylipins are more strongly related to IL-6/IL-8 or are predictors of outcomes or potential therapeutic response.

While our study identified relevant oxylipins associated with disease severity and etiology of ARDS, it has several limitations. As a novel exploratory analysis, it was not powered to link ARDS mortality to lipid species. In addition, we did not distinguish between pneumonia etiologies (i.e. viral or bacterial). Given the differences in oxylipins seen in viral pneumonias (such as COVID-19)(11) and bacterial pneumonia (36), this may have impacted differences between etiologies. Further, the risk of ARDS due to COVID-19 changed over the course of our sample collection, making it challenging to compare the effects as new viral variants emerged and vaccination and prior infections modified the host response (37). To partially account for this, our adjusted model (not shown here) included COVID-19 status, which was not associated with significant differences in PUFAs/oxylipins. Another limitation is that we measured plasma PUFAs, oxylipins, and markers of inflammation. Analysis of bronchoalveolar fluid may uncover additional insight.

Overall, our study adds to literature surrounding n-3 and n-6 PUFAs and their role in ARDS, identifying differences that may play a mechanistic role in ARDS. Identifying these distinct oxylipins which correlate with disease severity, inflammation, and ARDS etiology, increases our understanding of heterogeneity in ARDS.

## Data Availability Statement

The data that support the findings of this study are available in the Materials and Methods, Results, and/or Supplemental Material of this article. Any additional data is available upon reviewer’s request from the corresponding author.

## Author Contributions

LA Leuenberger and KM Gowdy conceived and designed the research study. RK Mallampalli, RM Tighe, KM Gowdy, LA Leuenberger, SR Shaikh contributed expertise to study design and data interpretation. JS Bednash, C Mikacenic, and ED Morrell provided experimental expertise and biorepository samples. E Schott and LA Leuenberger performed research experiments. C Richardson and LA Leuenberger performed statistical analyses. LA Leuenberger, E Schott, JC Horowitz, RM Tighe, RK Mallampalli, SR Shaikh, LA Fussner and KM Gowdy analyzed and interpreted the data. All authors contributed to drafting and revising the manuscript.

## Acknowledgements

The authors would like to thank Dr. Krishanarao Maddipati and the Wayne State Lipidomics Core for their expertise. We also wish to thank the patients who participated in these biorepositories, as well as their families and surrogates who served as patient advocates and decision makers and selflessly chose to participate in clinical research. Furthermore, we’d like to thank the investigators, clinical research coordinators and staff for their hard work collecting samples and data.

## References

1. Thompson BT, Chambers RC, Liu KD. Acute Respiratory Distress Syndrome. N Engl J Med. 2017;377(19):1904–5.

2. Calfee CS, Delucchi K, Parsons PE, Thompson BT, Ware LB, Matthay MA, et al. Subphenotypes in acute respiratory distress syndrome: latent class analysis of data from two randomised controlled trials. Lancet Respir Med. 2014;2(8):611–20.

3. Duvall MG, Levy BD. DHA- and EPA-derived resolvins, protectins, and maresins in airway inflammation. Eur J Pharmacol. 2016;785:144–55.

4. Stapleton RD, Martin TR, Weiss NS, Crowley JJ, Gundel SJ, Nathens AB, et al. A phase II randomized placebo-controlled trial of omega-3 fatty acids for the treatment of acute lung injury. Crit Care Med. 2011;39(7):1655–62.

5. Rice TW, Wheeler AP, Thompson BT, deBoisblanc BP, Steingrub J, Rock P, et al. Enteral omega-3 fatty acid, gamma-linolenic acid, and antioxidant supplementation in acute lung injury. JAMA. 2011;306(14):1574–81.

6. Fisk HL, Shaikh SR. Emerging mechanisms of organ crosstalk: The role of oxylipins. Nutr Bull. 2025;50(1):12–29.

7. Ekström S, Sdona E, Klevebro S, Hallberg J, Georgelis A, Kull I, et al. Dietary intake and plasma concentrations of PUFAs in childhood and adolescence in relation to asthma and lung function up to adulthood. Am J Clin Nutr. 2022;115(3):886–96.

8. Giudetti AM, Cagnazzo R. Beneficial effects of n-3 PUFA on chronic airway inflammatory diseases. Prostaglandins Other Lipid Mediat. 2012;99(3-4):57–67.

9. Kotlyarov S, Kotlyarova A. Molecular Mechanisms of Lipid Metabolism Disorders in Infectious Exacerbations of Chronic Obstructive Pulmonary Disease. Int J Mol Sci. 2021;22(14).

10. Velotti F, Costantini L, Merendino N. Omega-3 Polyunsaturated Fatty Acids (n-3 PUFAs) for Immunomodulation in COVID-19 Related Acute Respiratory Distress Syndrome (ARDS). J Clin Med. 2022;12(1).

11. Schwarz B, Sharma L, Roberts L, Peng X, Bermejo S, Leighton I, et al. Cutting Edge: Severe SARS-CoV-2 Infection in Humans Is Defined by a Shift in the Serum Lipidome, Resulting in Dysregulation of Eicosanoid Immune Mediators. J Immunol. 2021;206(2):329–34.

12. Chen H, Wang S, Zhao Y, Luo Y, Tong H, Su L. Correlation analysis of omega-3 fatty acids and mortality of sepsis and sepsis-induced ARDS in adults: data from previous randomized controlled trials. Nutr J. 2018;17(1):57.

13. Singer M, Deutschman CS, Seymour CW, Shankar-Hari M, Annane D, Bauer M, et al. The Third International Consensus Definitions for Sepsis and Septic Shock (Sepsis-3). JAMA. 2016;315(8):801–10.

14. Matthay MA, Arabi Y, Arroliga AC, Bernard G, Bersten AD, Brochard LJ, et al. A New Global Definition of Acute Respiratory Distress Syndrome. Am J Respir Crit Care Med. 2024;209(1):37–47.

15. Maddipati KR, Romero R, Chaiworapongsa T, Zhou SL, Xu Z, Tarca AL, et al. Eicosanomic profiling reveals dominance of the epoxygenase pathway in human amniotic fluid at term in spontaneous labor. FASEB J. 2014;28(11):4835–46.

16. Ranieri VM, Rubenfeld GD, Thompson BT, Ferguson ND, Caldwell E, Fan E, et al. Acute respiratory distress syndrome: the Berlin Definition. JAMA. 2012;307(23):2526–33.

17. Shah JS, Rai SN, DeFilippis AP, Hill BG, Bhatnagar A, Brock GN. Distribution based nearest neighbor imputation for truncated high dimensional data with applications to pre-clinical and clinical metabolomics studies. BMC Bioinformatics. 2017;18(1):114.

18. Benjamini Y, Hochberg Y. Controlling the False Discovery Rate: A Practical and Powerful Approach to Multiple Testing. Journal of the Royal Statistical Society Series B (Methodological). 1995;57(1):289–300.

19. Bountziouka V, Nelson CP, Codd V, Samani NJ. Higher dietary n - 3 PUFA and fiber intake are associated with longer leukocyte telomere length: Evidence from a substitution model analysis in the UK Biobank. Nutr Res. 2025;142:63–75.

20. Rotarescu RD, Mathur M, Green MR, Anderson GH, Metherel AH. Heart DHA turnover is faster in female compared to male ALA- and EPA-fed mice. J Lipid Res. 2025;66(10):100897.

21. Sahebkar A, Simental-Mendía LE, Pedone C, Ferretti G, Nachtigal P, Bo S, et al. Statin therapy and plasma free fatty acids: a systematic review and meta-analysis of controlled clinical trials. Br J Clin Pharmacol. 2016;81(5):807–18.

22. Bayram S, Kızıltan G. The Role of Omega- 3 Polyunsaturated Fatty Acids in Diabetes Mellitus Management: A Narrative Review. Curr Nutr Rep. 2024;13(3):527–51.

23. Zhang Y, Sun Y, Yu Q, Song S, Brenna JT, Shen Y, et al. Higher ratio of plasma omega-6/omega-3 fatty acids is associated with greater risk of all-cause, cancer, and cardiovascular mortality: A population-based cohort study in UK Biobank. Elife. 2024;12.

24. Seki H, Fukunaga K, Arita M, Arai H, Nakanishi H, Taguchi R, et al. The anti-inflammatory and proresolving mediator resolvin E1 protects mice from bacterial pneumonia and acute lung injury. J Immunol. 2010;184(2):836–43.

25. Kumar KV, Rao SM, Gayani R, Mohan IK, Naidu MU. Oxidant stress and essential fatty acids in patients with risk and established ARDS. Clin Chim Acta. 2000;298(1-2):111–20.

26. Luo J, Zhang WY, Li H, Zhang PH, Tian C, Wu CH, et al. Pro-Resolving Mediator Resolvin E1 Restores Alveolar Fluid Clearance in Acute Respiratory Distress Syndrome. Shock. 2022;57(4):565–75.

27. Nirengi S, Buck B, Das D, Peres Valgas da Silva C, Calyeca J, Baer LA, et al. 12,13-diHOME protects against the age-related decline in cardiovascular function via attenuation of CaMKII. Nat Commun. 2025;16(1):7088.

28. Insuela DBR, Ferrero MR, Coutinho DS, Martins MA, Carvalho VF. Could Arachidonic Acid-Derived Pro-Resolving Mediators Be a New Therapeutic Strategy for Asthma Therapy? Front Immunol. 2020;11:580598.

29. Li J, Guo C, Wu J. 15-Deoxy-Δ-. PPAR Res. 2019;2019:7242030.

30. Ashina M, Fujioka K, Nishida K, Okubo S, Ikuta T, Shinohara M, et al. Recombinant human thrombomodulin attenuated sepsis severity in a non-surgical preterm mouse model. Sci Rep. 2020;10(1):333.

31. Serhan CN, Chiang N, Van Dyke TE. Resolving inflammation: dual anti-inflammatory and pro-resolution lipid mediators. Nat Rev Immunol. 2008;8(5):349–61.

32. McAuley DF, Laffey JG, O’Kane CM, Perkins GD, Mullan B, Trinder TJ, et al. Simvastatin in the acute respiratory distress syndrome. N Engl J Med. 2014;371(18):1695–703.

33. Truwit JD, Bernard GR, Steingrub J, Matthay MA, Liu KD, Albertson TE, et al. Rosuvastatin for sepsis-associated acute respiratory distress syndrome. N Engl J Med. 2014;370(23):2191–200.

34. Yaeger MJ, Ngatikaura T, Zecchino N, Lovins HB, Schott E, Cochran SJ, et al. Docosahexaenoic Acid and Its Metabolites Protects Against Ozone-induced Pulmonary Inflammation. Am J Respir Cell Mol Biol. 2025.

35. Keenan AH, Pedersen TL, Fillaus K, Larson MK, Shearer GC, Newman JW. Basal omega-3 fatty acid status affects fatty acid and oxylipin responses to high-dose n3-HUFA in healthy volunteers. J Lipid Res. 2012;53(8):1662–9.

36. Georgakopoulou VE, Dodos K, Pitiriga VC. Role of Lipidomics in Respiratory Tract Infections: A Systematic Review of Emerging Evidence. Microorganisms. 2025;13(9).

37. El Zawily A, Eckert S, Adajar R, Wagih N, Elmaidomy AH, Helmy AM, et al. Comprehensive review on COVID-19: etiology, pathogenicity, and treatment. Front Med (Lausanne). 2025;12:1569013.

