## Supplemental Tables for "Heterogeneous causes of acute respiratory distress syndrome correlate with distinct peripheral polyunsaturated fatty acid metabolites"

### Supplemental Table 1: List of Oxylipins measured by LC-MS/MS

**Prostaglandins (PG):** PGE1, 13,14-dh-PGE1, 13,14-dh-15k-PGE1, 15-keto-PGE1, bicyclo-PGE1, 2,3dinor-PGE1, 6-keto-PGE1, PGE2, 15-keto-PGE2, 13,14-dh-15k-PGE2, bicyclo-PGE2, PGE3, PGJ2, d12-PGJ2, 15d-d12,1-PGJ2, PGF1a, 15-keto-PGF2a, 19R(OH)PGF2a, 20(OH)-PGF2a, 9-iso-PGF2, 11b-PGF2, 6,16, diketo-PGFa, PGD2, PGD3, PGF2a, PGF3a, 6-keto PGF1a,

**Thromboxanes (TX):** TXB2, TXB3, 2,3-dinor-TXB2, 11-dh-2,3-dinor-TXB2, 11-dh-TXB3

**Leukotrienes (LT):** LTB4, 20-hydroxy LTB4, LTB5, 20-COOH LTB4, 18-carboxy dinor LTB4

**Di-Hydroxyeicostetraenoic acid (DiHETE)** : 5(S),12(S)-DiHETE, 8(S)_15(S)-DiHETE 5(S)_6(S)-DiHETE, 5(S)_15(S)-DiHETE

**Hydroxy octadecadienoic acid (HODE)**: 9-HODE, 13-HODE

**Hydroxyoctadecatrienoic acid (HOTrE):** 9-HOTrE, 13-HOTrE

**Hydroxyeicosadienoic acid (HEDE):** 11-HEDE, 15-HEDE

**Hydroxyeicosatrienoic acid (HETrE):** 8-HETrE, 5-HETrE, 5,6-diHETrE, 8,9-diHETrE, 11,12-diHETrE,14,15 -diHETrE

**Hydroxyeicosatetraenoic Acid (HETE):** 5-HETE, 12-HETE, 9-HETE, 11-HETE, 12-HETE, 15-HETE, 20-HETE, tetranor-12-HETE

**Hydroxyicosapentaenoic Acids (HEPE)**: 5-HEPE, 8-HEPE, 9-HEPE, 11-HEPE, 12-HEPE, 15-HEPE, 18-HEPE; 5(S),15(S)-DiHEPE

**Hydroxydocosahexanoic acid (HDoHE):** 4-HDoHE, 7-HDoHE, 8-HDoHE, 10-HDoHE, 11-HDoHE, 13-HDoHE, 14-HDoHE, 16-HDoHE, 17-HDoHE, 20-HDoHE

**Epoxyeicosatrienoic acid (EpETrE)**:14,15-EpETrE

**Epoxyeicosatetraenoic acid (EpETE):** 14,15-EpETE, 17, 18-EpETE

**Epoxy docosapentaenoic acid (EpDPE):** 13,14-EpDPE, 19,20-EpDPE

**Dihydroxydocosapentaenoic acid (DiHDoPE):** 19,20-DiHDoPE

**Dihydroxyoctadecenoic acid (diHOME):** 9,10-DiHOME, 12,13-diHOME

**Oxo-octadecadienoic acid (oxoODE)**: 9-oxoODE, 13-oxoODE

**Oxo-octadecatrienoic acid (oxoOTrE):** 9-OxoOTrE

**Oxo-eicosatetraenoic acid (OxoETE):** 12-oxoETE, 15-oxoETE

**Oxo- eicosadienoic acid (OxoEDE):** 15-oxoEDE

**Lipoxins (LX):** LXA4, 15-epi LXA4, LXA5, LXB4,

**Resolvins (Rv): RvD1, Aspirin Triggered (AT)-RvD1,** RvD2, **R**vD3, AT-RvD3, RvD4, AT-RvD4, RvD5, RvD6, 8-oxo-RvD1, RvE1, RvE2,

**Protectin (PD):** PD1, AT-PD1, 22(OH)PD1

**Maresin (Mar):** Maresin1, Maresin2, MaR1(n-3 DPA)

**Supplemental Table 2**: **Oxylipin levels are not associated with mortality in Acute Respiratory Distress Syndrome (ARDS)**

| | Survived | | Died | |
|---|---|---|---|---|
| **Lipid Metabolite** | **avg** | **std** | **avg** | **std** |
| **13_14-dh-PGE1** | 1.588776 | 0.900733 | 1.558857 | 0.872993 |
| **13_14-dh-15k-PGE1** | 2.537647 | 1.629112 | 1.530769 | 0.987256 |
| **PGE1** | 2.758966 | 1.897595 | 3.205 | 1.413547 |
| **15(R)-PGE1** | 1.2425 | 0.658998 | 1.261667 | 0.604332 |
| **15-keto PGE1** | 0.598261 | 0.335508 | 0.516538 | 0.206416 |
| **Bicyclo PGE1** | 0.837714 | 0.468904 | 0.993103 | 0.976916 |
| **2_3-dinor PGE1** | 0.951429 | 0.89219 | 9.731765 | 21.09492 |
| **PGE2** | 0.958571 | 0.402841 | 0.989 | 0.468981 |
| **15-keto PGE2** | 1.009143 | 0.401617 | 1.075217 | 0.47216 |
| **13_14-dh-15k-PGE2** | 0.763333 | 0.595335 | 0.6575 | 0.438496 |
| **Bicyclo PGE2** | 2.473125 | 1.56271 | 2.058333 | 1.416685 |
| **tetranor PGEM** | 0.51 | 0.339411 | 0.233333 | 0.020817 |
| **PGE3** | 50.93 | 88.84319 | 103.1717 | 328.8162 |
| **PGD2** | 1.758182 | 1.140156 | 2.143333 | 1.526233 |
| **PGJ2** | 0.612143 | 0.278324 | 0.554 | 0.253605 |
| **D12-PGJ2** | 2.422941 | 2.112738 | 2.900263 | 4.798291 |
| **15d-D12_14-PGJ2** | 3.374423 | 7.422262 | 1.951622 | 3.993569 |
| **13_14-dh-15k-PGD2** | 1.111111 | 0.365289 | 0.72 | 0.417313 |
| **PGD3** | 1.1 | 0.353553 | 1.091429 | 0.473618 |
| **15d-D12_14-PGJ3** | 27.94346 | 49.67712 | 19.10105 | 38.24988 |
| **PGF1a** | 0.791429 | 0.690183 | 0.703846 | 0.431152 |
| **PGF2a** | 1.042222 | 0.778119 | 1.306 | 1.049824 |
| **15-keto PGF2a** | 0.4252 | 0.234612 | 0.556111 | 0.483628 |
| **19(R)-OH PGF2a & 20-OH PGF2a** | 234.7858 | 407.0705 | 271.8645 | 690.9398 |
| **PGF3a** | 2.7225 | 0.270231 | 2.19 | 0.713723 |
| **8-isoPGF2a & 11bPGF2a** | 0.955 | 0.570506 | 0.6592 | 0.406867 |
| **6-keto PGE1** | 0.204615 | 0.171866 | 1.219 | 3.403754 |
| **6_15-diketo PGFa** | 0.765385 | 0.78208 | 0.500385 | 0.312723 |
| **TXB2** | 1.166 | 1.933786 | 0.736667 | 0.291319 |
| **11-dh-TXB2** | 0.217143 | 0.209023 | 0.207 | 0.141661 |
| **2_3-dinor TXB2** | 0.916154 | 0.613046 | 0.74 | 0.607242 |
| **11-dh-2_3-dinor TXB2** | 2.517333 | 0.922983 | 2.415938 | 1.120092 |
| **TXB3** | 0.238333 | 0.266414 | 0.13125 | 0.089672 |
| **11-dh TXB3** | 0.600667 | 0.349274 | 0.678824 | 0.761453 |
| **LTB4** | 0.074231 | 0.04884 | 0.12 | 0.128695 |
| **12-OxoLTB4** | 0.031667 | 0.02137 | 0.017 | 0.009487 |
| **20-hydroxy LTB4** | 0.02625 | 0.015059 | 0.0275 | 0.012583 |
| **20-COOH LTB4** | 0.068 | 0.095778 | 0.035 | 0.049497 |

| | | | | |
|---|---|---|---|---|
| **18-carboxy dinor LTB4** | 1.05 | 1.244508 | 0.43 | ND |
| **LTB5** | 0.046458 | 0.031045 | 0.084118 | 0.114313 |
| **5(S)_6(S)-DiHETE** | 0.01 | 0 | 0.01 | 0.01 |
| **5(S)_12(S)-DiHETE** | 0.027917 | 0.01744 | 0.026471 | 0.023964 |
| **5(S)_15(S)-DiHETE** | 0.014444 | 0.007048 | 0.016 | 0.01075 |
| **8(S)_15(S)-DiHETE** | 0.04625 | 0.02143 | 0.099545 | 0.085717 |
| **5(S)_15(S)-DiHEPE** | 0.111 | 0.075537 | 0.220435 | 0.286142 |
| **9-HODE** | 59.47432 | 73.42975 | 40.40233 | 28.20287 |
| **13-HODE** | 257.2667 | 253.6035 | 308.4282 | 270.4264 |
| **9-HOTrE** | 6.240962 | 11.79942 | 6.065 | 8.621695 |
| **13-HOTrE** | 6.555682 | 8.684324 | 6.486552 | 6.827536 |
| **11-HEDE** | 0.686154 | 0.80496 | 0.88 | 1.103853 |
| **15-HEDE** | 0.375577 | 0.403827 | 0.552895 | 0.819518 |
| **8-HETrE** | 0.515962 | 0.34333 | 0.546053 | 0.392604 |
| **5-HETrE** | 0.189808 | 0.280829 | 0.194286 | 0.232964 |
| **5-HETE** | 2.320192 | 1.485556 | 2.561842 | 1.612781 |
| **8-HETE** | 1.393654 | 0.88673 | 1.677632 | 1.25858 |
| **9-HETE** | 0.6725 | 0.913053 | 1.008333 | 1.346697 |
| **11-HETE** | 3.636731 | 2.126685 | 5.049737 | 4.036329 |
| **12-HETE** | 25.3875 | 41.80575 | 34.13395 | 41.81988 |
| **15-HETE** | 4.1292 | 3.946202 | 4.797368 | 3.119859 |
| **20-HETE** | 1.52 | 1.259281 | 1.909211 | 1.410981 |
| **tetranor 12-HETE** | 0.469038 | 0.676662 | 0.517895 | 0.678564 |
| **12-HHTrE** | 0.34625 | 0.122584 | 0.37 | 0.375979 |
| **5-HEPE** | 0.47 | 0.74 | 0.52 | 0.85 |
| **8-HEPE** | 0.35 | 0.60 | 0.48 | 1.19 |
| **9-HEPE** | 0.39 | 0.67 | 2.06 | 5.03 |
| **11-HEPE** | 0.74 | 1.34 | 1.24 | 3.82 |
| **12-HEPE** | 1.99 | 4.14 | 7.75 | 36.38 |
| **15-HEPE** | 0.73 | 0.89 | 1.82 | 4.15 |
| **18-HEPE** | 0.65 | 1.05 | 0.95 | 2.62 |
| **4-HDoHE** | 0.78 | 0.52 | 0.92 | 0.86 |
| **7-HDoHE** | 0.26 | 0.18 | 0.30 | 0.38 |
| **8-HDoHE** | 0.38 | 0.32 | 0.47 | 0.73 |
| **10-HDoHE** | 0.59 | 0.59 | 0.88 | 1.88 |
| **11-HDoHE** | 0.64 | 0.63 | 1.12 | 2.63 |
| **13-HDoHE** | 0.61 | 0.68 | 0.99 | 2.39 |
| **14-HDoHE** | 2.03 | 3.53 | 4.21 | 12.83 |
| **16-HDoHE** | 1.21 | 1.19 | 1.83 | 3.28 |
| **17-HDoHE** | 1.15 | 1.12 | 1.28 | 1.27 |
| **20-HDoHE** | 2.11 | 2.36 | 5.31 | 14.94 |
| **9(10)-EpOME** | 24.64 | 17.02 | 18.91 | 13.31 |
| **5(6)-EpETrE** | 0.03 | 0.01 | 0.02 | 0.01 |

| | | | | |
|---|---|---|---|---|
| **8(9)-EpETrE** | 0.09 | 0.07 | 0.14 | 0.09 |
| **14(15)-EpETrE** | 0.19 | 0.11 | 0.26 | 0.21 |
| **8(9)-EpETE** | 0.03 | 0.02 | 0.03 | ND |
| **11(12)-EpETE** | 0.08 | 0.06 | 0.21 | 0.44 |
| **14(15)-EpETE** | 0.11 | 0.09 | 0.13 | 0.08 |
| **17(18)-EpETE** | 0.17 | 0.26 | 0.30 | 0.87 |
| **7(8)-EpDPE** | 0.02 | 0.01 | 0.03 | 0.01 |
| **13(14)-EpDPE** | 0.15 | 0.12 | 0.17 | 0.15 |
| **16(17)-EpDPE** | 0.29 | 0.24 | 0.20 | 0.09 |
| **19(20)-EpDPE** | 0.45 | 0.28 | 0.65 | 1.10 |
| **9_10-DiHOME** | 2.93 | 3.63 | 2.57 | 4.23 |
| **12_13-DiHOME** | 9.20 | 13.59 | 8.17 | 15.02 |
| **5_6-DiHETrE** | 0.06 | 0.04 | 0.04 | 0.02 |
| **8_9-DiHETrE** | 0.07 | 0.07 | 0.09 | 0.11 |
| **11_12-DiHETrE** | 0.38 | 0.20 | 0.47 | 0.48 |
| **14_15-DiHETrE** | 0.58 | 0.38 | 0.89 | 1.45 |
| **5_6-DiHETE(EPA)** | 0.01 | 0.01 | 0.01 | 0.01 |
| **19_20-DiHDoPE** | 0.54 | 0.45 | 0.66 | 0.68 |
| **9-OxoODE** | 209.67 | 144.10 | 231.79 | 179.43 |
| **13-OxoODE** | 134.40 | 143.02 | 163.14 | 172.71 |
| **9-OxoOTrE** | 7.69 | 7.35 | 6.89 | 6.69 |
| **15-OxoEDE** | 0.55 | 0.86 | 0.57 | 0.49 |
| **5-oxoETE** | 0.72 | 0.33 | 1.11 | 0.84 |
| **12-OxoETE** | 0.65 | 0.31 | 1.06 | 0.76 |
| **15-OxoETE** | 1.31 | 1.11 | 1.37 | 1.05 |
| **LXA4** | 0.24 | 0.15 | 0.23 | 0.13 |
| **15-epi LXA4** | 0.21 | 0.14 | 0.26 | 0.15 |
| **15-oxo LXA4** | 0.15 | 0.10 | 0.15 | 0.10 |
| **LXA5** | 0.12 | 0.06 | 0.15 | 0.08 |
| **LXB4** | 5.82 | 5.28 | 3.73 | 3.36 |
| **RvD1 & AT-RvD1** | 0.10 | 0.06 | 0.19 | 0.17 |
| **RvD3** | 0.08 | 0.04 | 0.07 | 0.06 |
| **AT-RvD3** | 0.04 | 0.02 | 0.06 | 0.04 |
| **RvD4** | 0.10 | 0.08 | 0.08 | 0.03 |
| **AT-RvD4** | 0.07 | 0.06 | 0.07 | 0.08 |
| **RvD5** | 0.08 | 0.02 | 0.17 | 0.19 |
| **RvD6** | 0.17 | 0.08 | 0.17 | 0.13 |
| **AT-RvD6** | 0.28 | 0.13 | 0.26 | 0.26 |
| **8-oxoRvD1** | 0.17 | 0.07 | 0.14 | 0.08 |
| **17-oxoRvD1** | 0.08 | 0.03 | 0.07 | 0.02 |
| **RvD5 (n-3_DPA)** | 0.20 | 0.19 | 0.23 | 0.12 |
| **RvE1** | 0.93 | 1.55 | 1.38 | 2.26 |
| **RvE2** | 0.04 | 0.03 | 0.01 | 0.01 |

| | | | | |
|---|---|---|---|---|
| **RvE3** | 0.01 | 0.01 | 0.01 | 0.00 |
| **PD1** | 0.01 | 0.01 | 0.02 | 0.03 |
| **AT-PD1** | 0.14 | 0.30 | 0.24 | 0.41 |
| **PD1(n-3_DPA)** | 0.01 | 0.01 | 0.02 | 0.01 |
| **22-OH-PD1** | 0.06 | 0.06 | 0.08 | 0.11 |
| **Maresin1** | 0.01 | 0.01 | 0.01 | 0.00 |
| **Maresin2** | 0.02 | 0.01 | 0.04 | 0.05 |
| **MaR1(n-3_DPA)** | 0.29 | 0.19 | 0.51 | 0.36 |

Metabolites measured by LC/MS-MS. Table shows average and standard deviation for concentration (ng/mL) of metabolites, which were found to not be significantly different when modeled by multivariate regression. Results were the same in both adjusted and unadjusted models. N=52 survived, 38 died.

**Supplemental Table 3: Multiple n-6 polyunsaturated fatty acids are increased in patients with ARDS secondary to Aspiration, Trauma and Other causes compared to ARDS secondary to Sepsis**

| PUFA | Outcome | Beta | 95% CI | *P*-value | *Q*-value |
|---|---|---|---|---|---|
| **ALA + GLA** | Sepsis | — | — | | |
| | Aspiration | 0.92 | 0.39, 1.4 | <0.001 | **0.009** |
| | Other | 0.88 | 0.06, 1.7 | 0.036 | 0.13 |
| | Pneumonia | 0.49 | -0.07, 1.0 | 0.085 | 0.2 |
| | Trauma | 1.4 | 0.77, 2.0 | <0.001 | **<0.001** |
| **SDA** | Sepsis | — | — | | |
| | Aspiration | -0.63 | -1.5, 0.20 | 0.13 | 0.3 |
| | Other | 0.45 | -0.83, 1.7 | 0.5 | 0.6 |
| | Pneumonia | -0.63 | -1.5, 0.24 | 0.2 | 0.3 |
| | Trauma | -0.33 | -1.3, 0.67 | 0.5 | 0.7 |
| **EPA** | Sepsis | — | — | | |
| | Aspiration | -0.46 | -1.1, 0.20 | 0.2 | 0.4 |
| | Other | -0.12 | -1.2, 0.91 | 0.8 | 0.9 |
| | Pneumonia | -0.39 | -1.1, 0.31 | 0.3 | 0.5 |
| | Trauma | -0.04 | -0.84, 0.76 | >0.9 | >0.9 |
| **n-3 DPA** | Sepsis | — | — | | |
| | Aspiration | -0.07 | -0.41, 0.27 | 0.7 | 0.8 |
| | Other | 0.21 | -0.31, 0.74 | 0.4 | 0.6 |
| | Pneumonia | 0.04 | -0.32, 0.39 | 0.8 | 0.9 |
| | Trauma | -0.22 | -0.63, 0.19 | 0.3 | 0.5 |
| **DHA** | Sepsis | — | — | | |
| | Aspiration | -0.24 | -0.60, 0.12 | 0.2 | 0.4 |
| | Other | 0.27 | -0.28, 0.82 | 0.3 | 0.5 |
| | Pneumonia | 0.02 | -0.35, 0.39 | >0.9 | >0.9 |
| | Trauma | 0.05 | -0.38, 0.48 | 0.8 | 0.9 |
| **LA** | Sepsis | — | — | | |
| | Aspiration | 0.37 | 0.14, 0.59 | 0.002 | **0.015** |
| | Other | 0.81 | 0.46, 1.2 | <0.001 | **<0.001** |
| | Pneumonia | 0.24 | 0.01, 0.48 | 0.043 | 0.15 |
| | Trauma | 0.63 | 0.36, 0.90 | <0.001 | **<0.001** |
| **DGLA** | Sepsis | — | — | | |
| | Aspiration | 0.19 | -0.23, 0.61 | 0.4 | 0.6 |
| | Other | 0.62 | -0.02, 1.3 | 0.058 | 0.2 |
| | Pneumonia | 0.43 | -0.01, 0.87 | 0.054 | 0.2 |
| | Trauma | 0.75 | 0.24, 1.3 | 0.004 | **0.023** |
| **AA** | Sepsis | — | — | | |
| | Aspiration | -0.07 | -0.35, 0.20 | 0.6 | 0.7 |
| | Other | 0.19 | -0.23, 0.62 | 0.4 | 0.6 |
| | Pneumonia | -0.02 | -0.31, 0.27 | 0.9 | >0.9 |
| | Trauma | -0.01 | -0.34, 0.32 | >0.9 | >0.9 |

| | | | | | |
|---|---|---|---|---|---|
| **n-6 DPA** | Sepsis | — | — | | |
| | Aspiration | 0.19 | -0.15, 0.53 | 0.3 | 0.5 |
| | Other | 0.68 | 0.15, 1.2 | 0.013 | **0.06** |
| | Pneumonia | 0.09 | -0.27, 0.45 | 0.6 | 0.7 |
| | Trauma | 0.19 | -0.22, 0.60 | 0.4 | 0.5 |

Polyunsaturated fatty acids (PUFAs) were measured by LC/MS-MS. Tables shows beta (estimate) and confidence intervals for PUFAs as compared concentration seen in sepsis, with the Benjamini-Hochberg false discovery rate set for a significance threshold of $Q<0.1$. Significant results in **bold.** N=90.

**Supplemental Table 4: Patients with ARDS secondary to Sepsis have differences in oxylipins compared to patients with other causes of ARDS.**

| Oxylipin | Outcome | Beta | 95% CI | *P*-value | *Q*-value |
|---|---|---|---|---|---|
| **13,14-dh-PGE** | Sepsis | — | — | | |
| | Aspiration | 0.45 | -0.26, 1.1 | 0.2 | 0.4 |
| | Other | -0.66 | -1.7, 0.43 | 0.2 | 0.4 |
| | Pneumonia | -0.05 | -0.78, 0.69 | 0.9 | >0.9 |
| | Trauma | 0.45 | -0.39, 1.3 | 0.3 | 0.5 |
| **13,14-dh-15k-PGE** | Sepsis | — | — | | |
| | Aspiration | 0.58 | 0.08, 1.1 | 0.024 | **0.086** |
| | Other | -0.29 | -1.1, 0.48 | 0.5 | 0.6 |
| | Pneumonia | 0.18 | -0.35, 0.70 | 0.5 | 0.7 |
| | Trauma | 0.52 | -0.09, 1.1 | 0.093 | 0.2 |
| **PGE1** | Sepsis | — | — | | |
| | Aspiration | 0.39 | -0.29, 1.1 | 0.3 | 0.5 |
| | Other | -0.44 | -1.5, 0.61 | 0.4 | 0.6 |
| | Pneumonia | 0.25 | -0.46, 0.96 | 0.5 | 0.7 |
| | Trauma | 0.37 | -0.45, 1.2 | 0.4 | 0.6 |
| **15-keto-PGE1** | Sepsis | — | — | | |
| | Aspiration | 0.68 | 0.16, 1.2 | 0.011 | **0.053** |
| | Other | -0.2 | -1.0, 0.60 | 0.6 | 0.8 |
| | Pneumonia | 0.71 | 0.17, 1.3 | 0.011 | **0.053** |
| | Trauma | 0.6 | -0.03, 1.2 | 0.061 | 0.2 |
| **bicyclo-PGE1** | Sepsis | — | — | | |
| | Aspiration | 0.57 | -0.01, 1.2 | 0.056 | 0.2 |
| | Other | 0.04 | -0.86, 0.94 | >0.9 | >0.9 |
| | Pneumonia | 0.3 | -0.30, 0.91 | 0.3 | 0.5 |
| | Trauma | 0.67 | -0.03, 1.4 | 0.059 | 0.2 |
| **2,3-dinor-PGE1** | Sepsis | — | — | | |
| | Aspiration | 2.1 | 1.2, 2.9 | <0.001 | **<0.001** |
| | Other | 0.43 | -0.85, 1.7 | 0.5 | 0.7 |
| | Pneumonia | 0.6 | -0.27, 1.5 | 0.2 | 0.4 |
| | Trauma | 1.5 | 0.47, 2.5 | 0.004 | **0.028** |
| **15-keto-PGE2** | Sepsis | — | — | | |
| | Aspiration | 0.18 | -0.13, 0.50 | 0.3 | 0.4 |
| | Other | 0.38 | -0.11, 0.87 | 0.13 | 0.3 |
| | Pneumonia | 0.27 | -0.06, 0.60 | 0.11 | 0.3 |
| | Trauma | 0.08 | -0.30, 0.46 | 0.7 | 0.8 |
| **13,14-dh-15k-PGE2** | Sepsis | — | — | | |
| | Aspiration | 0.94 | 0.43, 1.5 | <0.001 | **0.006** |
| | Other | 0.09 | -0.70, 0.88 | 0.8 | >0.9 |
| | Pneumonia | 0.49 | -0.04, 1.0 | 0.072 | 0.2 |
| | Trauma | 0.99 | 0.38, 1.6 | 0.002 | **0.017** |
| **bicyclo-PGE2** | Sepsis | — | — | | |
| | Aspiration | 0.78 | 0.44, 1.1 | <0.001 | **<0.001** |
| | Other | 0.44 | -0.09, 0.97 | 0.1 | 0.2 |
| | Pneumonia | 0.57 | 0.22, 0.93 | 0.002 | **0.018** |
| | Trauma | 0.63 | 0.22, 1.0 | 0.003 | **0.024** |

| | | | | | |
|---|---|---|---|---|---|
| **PGE3** | Sepsis | — | — | | |
| | Aspiration | 0.7 | -0.52, 1.9 | 0.3 | 0.5 |
| | Other | 1.3 | -0.60, 3.2 | 0.2 | 0.4 |
| | Pneumonia | 0.94 | -0.33, 2.2 | 0.15 | 0.3 |
| | Trauma | 1.3 | -0.15, 2.8 | 0.079 | 0.2 |
| **PGJ2** | Sepsis | — | — | | |
| | Aspiration | 0.45 | 0.12, 0.78 | 0.009 | **0.045** |
| | Other | 0.33 | -0.18, 0.84 | 0.2 | 0.4 |
| | Pneumonia | 0.22 | -0.13, 0.56 | 0.2 | 0.4 |
| | Trauma | 0.21 | -0.19, 0.61 | 0.3 | 0.5 |
| **d12-PGJ2** | Sepsis | — | — | | |
| | Aspiration | 1.5 | 0.74, 2.2 | <0.001 | **0.003** |
| | Other | 0.07 | -1.1, 1.2 | >0.9 | >0.9 |
| | Pneumonia | 1 | 0.28, 1.8 | 0.008 | **0.043** |
| | Trauma | 1.4 | 0.52, 2.3 | 0.002 | **0.018** |
| **15d-d12,14-PGJ2** | Sepsis | — | — | | |
| | Aspiration | 2.4 | 1.4, 3.4 | <0.001 | **<0.001** |
| | Other | 1.9 | 0.38, 3.5 | 0.015 | **0.067** |
| | Pneumonia | 0.75 | -0.31, 1.8 | 0.2 | 0.3 |
| | Trauma | 1.8 | 0.60, 3.0 | 0.004 | **0.026** |
| **15d-d12,14-PGJ3** | Sepsis | — | — | | |
| | Aspiration | 2.2 | 1.3, 3.1 | <0.001 | **<0.001** |
| | Other | 1.7 | 0.33, 3.2 | 0.017 | **0.07** |
| | Pneumonia | 1 | 0.07, 2.0 | 0.036 | 0.12 |
| | Trauma | 2.5 | 1.3, 3.6 | <0.001 | **<0.001** |
| **PGF1a** | Sepsis | — | — | | |
| | Aspiration | 0.93 | 0.54, 1.3 | <0.001 | **<0.001** |
| | Other | 0.5 | -0.11, 1.1 | 0.11 | 0.3 |
| | Pneumonia | 0.51 | 0.09, 0.92 | 0.017 | **0.07** |
| | Trauma | 1.3 | 0.84, 1.8 | <0.001 | **<0.001** |
| **15-keto-PGF2a** | Sepsis | — | — | | |
| | Aspiration | 0.38 | -0.18, 0.93 | 0.2 | 0.4 |
| | Other | -0.44 | -1.3, 0.41 | 0.3 | 0.5 |
| | Pneumonia | 0.28 | -0.29, 0.86 | 0.3 | 0.5 |
| | Trauma | 0.75 | 0.09, 1.4 | 0.027 | **0.095** |
| **19-OH-PGF2a** | Sepsis | — | — | | |
| | Aspiration | 1.8 | 0.05, 3.6 | 0.044 | 0.14 |
| | Other | 4.1 | 1.4, 6.9 | 0.003 | **0.024** |
| | Pneumonia | 1 | -0.81, 2.9 | 0.3 | 0.5 |
| | Trauma | 3.5 | 1.3, 5.6 | 0.002 | **0.017** |
| **8-isoPGF2a** | Sepsis | — | — | | |
| | Aspiration | 1 | 0.61, 1.4 | <0.001 | **<0.001** |
| | Other | 1.1 | 0.43, 1.7 | 0.001 | **0.013** |
| | Pneumonia | 0.63 | 0.21, 1.1 | 0.004 | **0.025** |
| | Trauma | 1.5 | 0.98, 1.9 | <0.001 | **<0.001** |
| **6-keto-PGE1** | Sepsis | — | — | | |
| | Aspiration | 1 | 0.22, 1.8 | 0.013 | **0.058** |
| | Other | -0.04 | -1.2, 1.2 | >0.9 | >0.9 |
| | Pneumonia | 0.39 | -0.43, 1.2 | 0.3 | 0.5 |
| | Trauma | 1.6 | 0.68, 2.6 | <0.001 | **0.011** |

| | | | | | |
|---|---|---|---|---|---|
| **6,15-diketo-PGFa** | Sepsis | — | — | | |
| | Aspiration | 1 | 0.54, 1.5 | <0.001 | **0.001** |
| | Other | 0.45 | -0.30, 1.2 | 0.2 | 0.4 |
| | Pneumonia | 0.05 | -0.46, 0.56 | 0.8 | >0.9 |
| | Trauma | 1.1 | 0.51, 1.7 | <0.001 | **0.006** |
| **2,3-dinor-TXB2** | Sepsis | — | — | | |
| | Aspiration | 0.25 | -0.09, 0.58 | 0.15 | 0.3 |
| | Other | 0.29 | -0.23, 0.80 | 0.3 | 0.5 |
| | Pneumonia | -0.08 | -0.43, 0.27 | 0.6 | 0.8 |
| | Trauma | 0.74 | 0.33, 1.1 | <0.001 | **0.007** |
| **11,dh-2,3-dinor-TXB2** | Sepsis | — | — | | |
| | Aspiration | 0.54 | 0.08, 1.0 | 0.021 | **0.082** |
| | Other | -0.61 | -1.3, 0.10 | 0.091 | 0.2 |
| | Pneumonia | 0.3 | -0.18, 0.78 | 0.2 | 0.4 |
| | Trauma | 0.34 | -0.21, 0.89 | 0.2 | 0.4 |
| **11-dh-TXB3** | Sepsis | — | — | | |
| | Aspiration | 1.1 | 0.66, 1.6 | <0.001 | **<0.001** |
| | Other | 0.09 | -0.63, 0.82 | 0.8 | >0.9 |
| | Pneumonia | 0.44 | -0.06, 0.93 | 0.081 | 0.2 |
| | Trauma | 1 | 0.46, 1.6 | <0.001 | **0.007** |
| **LTB4** | Sepsis | — | — | | |
| | Aspiration | 0.24 | -0.25, 0.74 | 0.3 | 0.5 |
| | Other | 0.07 | -0.69, 0.83 | 0.9 | >0.9 |
| | Pneumonia | 0.18 | -0.33, 0.70 | 0.5 | 0.7 |
| | Trauma | -0.59 | -1.2, 0.00 | 0.051 | 0.2 |
| **LTB5** | Sepsis | — | — | | |
| | Aspiration | 0.04 | -0.59, 0.67 | >0.9 | >0.9 |
| | Other | 1.1 | 0.17, 2.1 | 0.022 | **0.082** |
| | Pneumonia | 0.02 | -0.63, 0.68 | >0.9 | >0.9 |
| | Trauma | -0.54 | -1.3, 0.21 | 0.2 | 0.3 |
| **5(S),12(S)-diHETE** | Sepsis | — | — | | |
| | Aspiration | -0.12 | -0.53, 0.29 | 0.6 | 0.7 |
| | Other | 0.4 | -0.22, 1.0 | 0.2 | 0.4 |
| | Pneumonia | -0.03 | -0.45, 0.40 | 0.9 | >0.9 |
| | Trauma | -0.24 | -0.73, 0.25 | 0.3 | 0.5 |
| **-5(S),15(S)-diHETE** | Sepsis | — | — | | |
| | Aspiration | 0.57 | 0.33, 0.81 | <0.001 | **<0.001** |
| | Other | 0.41 | 0.04, 0.79 | 0.029 | 0.1 |
| | Pneumonia | 0.05 | -0.20, 0.30 | 0.7 | 0.8 |
| | Trauma | 0.41 | 0.12, 0.69 | 0.006 | **0.036** |
| **8(S),15(S)-diHETE** | Sepsis | — | — | | |
| | Aspiration | 0.16 | -0.35, 0.67 | 0.5 | 0.7 |
| | Other | 0.67 | -0.12, 1.5 | 0.094 | 0.2 |
| | Pneumonia | 0.26 | -0.27, 0.80 | 0.3 | 0.5 |
| | Trauma | -0.76 | -1.4, -0.15 | 0.015 | **0.067** |
| **5(S),15(S)-diHEPE** | Sepsis | — | — | | |
| | Aspiration | 0.07 | -0.43, 0.58 | 0.8 | 0.9 |
| | Other | 1.2 | 0.40, 2.0 | 0.003 | **0.024** |
| | Pneumonia | 0.26 | -0.27, 0.79 | 0.3 | 0.5 |
| | Trauma | -0.45 | -1.1, 0.16 | 0.15 | 0.3 |

| | | | | | |
|---|---|---|---|---|---|
| **9-HODE** | Sepsis | — | — | | |
| | Aspiration | 0.83 | 0.21, 1.4 | 0.009 | **0.045** |
| | Other | 0.64 | -0.31, 1.6 | 0.2 | 0.4 |
| | Pneumonia | 0.56 | -0.09, 1.2 | 0.089 | 0.2 |
| | Trauma | -0.09 | -0.83, 0.65 | 0.8 | >0.9 |
| **13-HODE** | Sepsis | — | — | | |
| | Aspiration | 0.77 | 0.14, 1.4 | 0.017 | **0.07** |
| | Other | 0.6 | -0.37, 1.6 | 0.2 | 0.4 |
| | Pneumonia | 0.6 | -0.05, 1.3 | 0.07 | 0.2 |
| | Trauma | -0.16 | -0.91, 0.60 | 0.7 | 0.8 |
| **9-HOTrE** | Sepsis | — | — | | |
| | Aspiration | 1.5 | 0.66, 2.3 | <0.001 | **0.007** |
| | Other | 1.2 | -0.10, 2.4 | 0.071 | 0.2 |
| | Pneumonia | 0.81 | -0.04, 1.7 | 0.063 | 0.2 |
| | Trauma | 0.44 | -0.54, 1.4 | 0.4 | 0.6 |
| **13-HOTrE** | Sepsis | — | — | | |
| | Aspiration | 1.3 | 0.61, 2.0 | <0.001 | **0.006** |
| | Other | 0.7 | -0.36, 1.8 | 0.2 | 0.4 |
| | Pneumonia | 0.81 | 0.09, 1.5 | 0.029 | 0.1 |
| | Trauma | 0.26 | -0.57, 1.1 | 0.5 | 0.7 |
| **11-HEDE** | Sepsis | — | — | | |
| | Aspiration | -0.13 | -0.77, 0.51 | 0.7 | 0.8 |
| | Other | 0.47 | -0.51, 1.4 | 0.3 | 0.5 |
| | Pneumonia | 0.13 | -0.53, 0.79 | 0.7 | 0.8 |
| | Trauma | -1.2 | -1.9, -0.41 | 0.003 | **0.024** |
| **15-HEDE** | Sepsis | — | — | | |
| | Aspiration | 0.26 | -0.49, 1.0 | 0.5 | 0.7 |
| | Other | 0.71 | -0.45, 1.9 | 0.2 | 0.4 |
| | Pneumonia | 0.55 | -0.24, 1.3 | 0.2 | 0.3 |
| | Trauma | -0.36 | -1.3, 0.54 | 0.4 | 0.6 |
| **8-HETrE** | Sepsis | — | — | | |
| | Aspiration | -0.22 | -0.70, 0.26 | 0.4 | 0.6 |
| | Other | 0.32 | -0.42, 1.1 | 0.4 | 0.6 |
| | Pneumonia | 0.26 | -0.25, 0.76 | 0.3 | 0.5 |
| | Trauma | -0.8 | -1.4, -0.22 | 0.007 | **0.038** |
| **5-HETrE** | Sepsis | — | — | | |
| | Aspiration | -0.01 | -0.65, 0.63 | >0.9 | >0.9 |
| | Other | 0.54 | -0.45, 1.5 | 0.3 | 0.5 |
| | Pneumonia | 0.12 | -0.55, 0.80 | 0.7 | 0.9 |
| | Trauma | -0.85 | -1.6, -0.08 | 0.031 | 0.1 |
| **5-HETE** | Sepsis | — | — | | |
| | Aspiration | -0.18 | -0.64, 0.27 | 0.4 | 0.6 |
| | Other | 0.26 | -0.45, 0.96 | 0.5 | 0.7 |
| | Pneumonia | 0.01 | -0.47, 0.49 | >0.9 | >0.9 |
| | Trauma | -0.99 | -1.5, -0.44 | <0.001 | **0.007** |
| **8-HETE** | Sepsis | — | — | | |
| | Aspiration | -0.32 | -0.77, 0.14 | 0.2 | 0.4 |
| | Other | 0.29 | -0.42, 0.99 | 0.4 | 0.6 |
| | Pneumonia | 0.06 | -0.42, 0.54 | 0.8 | >0.9 |
| | Trauma | -0.86 | -1.4, -0.31 | 0.003 | **0.022** |

| | | | | | |
|---|---|---|---|---|---|
| **9-HETE** | Sepsis | — | — | | |
| | Aspiration | 0.11 | -0.66, 0.87 | 0.8 | >0.9 |
| | Other | -0.13 | -1.3, 1.0 | 0.8 | >0.9 |
| | Pneumonia | 0.14 | -0.66, 0.93 | 0.7 | 0.9 |
| | Trauma | -0.87 | -1.8, 0.05 | 0.062 | 0.2 |
| **11-HETE** | Sepsis | — | — | | |
| | Aspiration | -0.46 | -0.97, 0.04 | 0.073 | 0.2 |
| | Other | 0.01 | -0.78, 0.79 | >0.9 | >0.9 |
| | Pneumonia | -0.03 | -0.56, 0.50 | >0.9 | >0.9 |
| | Trauma | -1.2 | -1.9, -0.64 | <0.001 | **0.002** |
| **12-HETE** | Sepsis | — | — | | |
| | Aspiration | -0.27 | -1.1, 0.56 | 0.5 | 0.7 |
| | Other | -0.57 | -1.9, 0.71 | 0.4 | 0.6 |
| | Pneumonia | -0.02 | -0.88, 0.85 | >0.9 | >0.9 |
| | Trauma | -1.5 | -2.5, -0.54 | 0.003 | **0.023** |
| **15-HETE** | Sepsis | — | — | | |
| | Aspiration | -0.53 | -1.1, 0.00 | 0.052 | 0.2 |
| | Other | -0.05 | -0.88, 0.77 | 0.9 | >0.9 |
| | Pneumonia | -0.2 | -0.76, 0.36 | 0.5 | 0.7 |
| | Trauma | -1 | -1.7, -0.37 | 0.002 | **0.019** |
| **20-HETE** | Sepsis | — | — | | |
| | Aspiration | -0.2 | -0.67, 0.28 | 0.4 | 0.6 |
| | Other | 0.19 | -0.55, 0.92 | 0.6 | 0.8 |
| | Pneumonia | -0.19 | -0.69, 0.31 | 0.4 | 0.6 |
| | Trauma | -0.92 | -1.5, -0.35 | 0.002 | **0.017** |
| **tetranor-12-HETE** | Sepsis | — | — | | |
| | Aspiration | 0.04 | -0.43, 0.51 | 0.9 | >0.9 |
| | Other | 0.5 | -0.22, 1.2 | 0.2 | 0.4 |
| | Pneumonia | -0.05 | -0.54, 0.44 | 0.8 | >0.9 |
| | Trauma | -0.23 | -0.79, 0.34 | 0.4 | 0.6 |
| **5-HEPE** | Sepsis | — | — | | |
| | Aspiration | -0.54 | -1.1, 0.05 | 0.072 | 0.2 |
| | Other | 0.03 | -0.89, 0.94 | >0.9 | >0.9 |
| | Pneumonia | -0.18 | -0.80, 0.45 | 0.6 | 0.7 |
| | Trauma | -0.77 | -1.5, -0.06 | 0.035 | 0.11 |
| **8-HEPE** | Sepsis | — | — | | |
| | Aspiration | -0.53 | -1.2, 0.18 | 0.14 | 0.3 |
| | Other | -0.35 | -1.4, 0.74 | 0.5 | 0.7 |
| | Pneumonia | -0.59 | -1.3, 0.15 | 0.12 | 0.3 |
| | Trauma | -0.92 | -1.8, -0.07 | 0.034 | 0.11 |
| **9-HEPE** | Sepsis | — | — | | |
| | Aspiration | -0.63 | -1.4, 0.19 | 0.13 | 0.3 |
| | Other | 0.12 | -1.1, 1.4 | 0.8 | >0.9 |
| | Pneumonia | -0.25 | -1.1, 0.60 | 0.6 | 0.7 |
| | Trauma | -0.91 | -1.9, 0.07 | 0.067 | 0.2 |
| **11-HEPE** | Sepsis | — | — | | |
| | Aspiration | -0.67 | -1.4, 0.06 | 0.071 | 0.2 |
| | Other | -0.07 | -1.2, 1.1 | >0.9 | >0.9 |
| | Pneumonia | -0.3 | -1.1, 0.46 | 0.4 | 0.6 |
| | Trauma | -1.1 | -1.9, -0.19 | 0.017 | **0.07** |

| | | | | | |
|---|---|---|---|---|---|
| **12-HEPE** | Sepsis | — | — | | |
| | Aspiration | -1.2 | -2.2, -0.17 | 0.023 | **0.086** |
| | Other | -1.1 | -2.7, 0.44 | 0.2 | 0.3 |
| | Pneumonia | -1.1 | -2.2, -0.03 | 0.044 | 0.14 |
| | Trauma | -1.8 | -3.0, -0.56 | 0.005 | **0.029** |
| **15-HEPE** | Sepsis | — | — | | |
| | Aspiration | -1.2 | -1.9, -0.49 | <0.001 | **0.011** |
| | Other | -0.62 | -1.7, 0.44 | 0.2 | 0.4 |
| | Pneumonia | -0.86 | -1.6, -0.15 | 0.018 | **0.072** |
| | Trauma | -2 | -2.8, -1.2 | <0.001 | **<0.001** |
| **18-HEPE** | Sepsis | — | — | | |
| | Aspiration | -0.6 | -1.3, 0.10 | 0.091 | 0.2 |
| | Other | -0.42 | -1.5, 0.65 | 0.4 | 0.6 |
| | Pneumonia | -0.43 | -1.2, 0.29 | 0.2 | 0.4 |
| | Trauma | -1.2 | -2.0, -0.34 | 0.006 | **0.036** |
| **4-HDoHE** | Sepsis | — | — | | |
| | Aspiration | -0.6 | -1.3, 0.10 | 0.091 | 0.2 |
| | Other | -0.42 | -1.5, 0.65 | 0.4 | 0.6 |
| | Pneumonia | -0.43 | -1.2, 0.29 | 0.2 | 0.4 |
| | Trauma | -1.2 | -2.0, -0.34 | 0.006 | **0.036** |
| **7-HDoHE** | Sepsis | — | — | | |
| | Aspiration | -0.39 | -0.91, 0.13 | 0.14 | 0.3 |
| | Other | 0.26 | -0.55, 1.1 | 0.5 | 0.7 |
| | Pneumonia | 0.01 | -0.54, 0.55 | >0.9 | >0.9 |
| | Trauma | -0.6 | -1.2, 0.03 | 0.062 | 0.2 |
| **8-HDoHE** | Sepsis | — | — | | |
| | Aspiration | -0.34 | -0.96, 0.28 | 0.3 | 0.5 |
| | Other | 0.29 | -0.67, 1.2 | 0.6 | 0.7 |
| | Pneumonia | 0.21 | -0.44, 0.86 | 0.5 | 0.7 |
| | Trauma | -0.4 | -1.1, 0.34 | 0.3 | 0.5 |
| **10-HDoHE** | Sepsis | — | — | | |
| | Aspiration | -0.41 | -1.1, 0.24 | 0.2 | 0.4 |
| | Other | -0.08 | -1.1, 0.93 | 0.9 | >0.9 |
| | Pneumonia | 0 | -0.68, 0.68 | >0.9 | >0.9 |
| | Trauma | -0.86 | -1.6, -0.07 | 0.032 | 0.11 |
| **11-HDoHE** | Sepsis | — | — | | |
| | Aspiration | -0.76 | -1.5, -0.07 | 0.032 | 0.11 |
| | Other | -0.21 | -1.3, 0.86 | 0.7 | 0.8 |
| | Pneumonia | -0.38 | -1.1, 0.35 | 0.3 | 0.5 |
| | Trauma | -1.4 | -2.2, -0.53 | 0.002 | **0.017** |
| **13-HDoHE** | Sepsis | — | — | | |
| | Aspiration | -0.6 | -1.3, 0.14 | 0.11 | 0.3 |
| | Other | 0.06 | -1.1, 1.2 | >0.9 | >0.9 |
| | Pneumonia | -0.13 | -0.89, 0.63 | 0.7 | 0.9 |
| | Trauma | -1.2 | -2.1, -0.30 | 0.009 | **0.045** |
| **14-HDoHE** | Sepsis | — | — | | |
| | Aspiration | -0.77 | -1.7, 0.13 | 0.091 | 0.2 |
| | Other | -1.1 | -2.5, 0.24 | 0.1 | 0.3 |
| | Pneumonia | -0.36 | -1.3, 0.57 | 0.4 | 0.6 |
| | Trauma | -1.7 | -2.8, -0.65 | 0.002 | **0.017** |

| | | | | | |
|---|---|---|---|---|---|
| **16-HDoHE** | Sepsis | — | — | | |
| | Aspiration | -0.36 | -1.0, 0.33 | 0.3 | 0.5 |
| | Other | 0.02 | -1.0, 1.1 | >0.9 | >0.9 |
| | Pneumonia | 0.04 | -0.68, 0.77 | >0.9 | >0.9 |
| | Trauma | -1 | -1.8, -0.19 | 0.017 | 0.07 |
| **17-HDoHE** | Sepsis | — | — | | |
| | Aspiration | -0.43 | -1.0, 0.18 | 0.2 | 0.3 |
| | Other | -0.03 | -0.97, 0.91 | >0.9 | >0.9 |
| | Pneumonia | -0.26 | -0.89, 0.38 | 0.4 | 0.6 |
| | Trauma | -1.4 | -2.1, -0.67 | <0.001 | **0.005** |
| **20-HDoHE** | Sepsis | — | — | | |
| | Aspiration | -0.27 | -1.1, 0.53 | 0.5 | 0.7 |
| | Other | 1.6 | 0.35, 2.8 | 0.013 | **0.058** |
| | Pneumonia | 0.06 | -0.78, 0.90 | 0.9 | >0.9 |
| | Trauma | 0.21 | -0.75, 1.2 | 0.7 | 0.8 |
| **8,9-EPETrE** | Sepsis | — | — | | |
| | Aspiration | -0.16 | -0.66, 0.34 | 0.5 | 0.7 |
| | Other | 0.1 | -0.68, 0.87 | 0.8 | >0.9 |
| | Pneumonia | 0.02 | -0.50, 0.54 | >0.9 | >0.9 |
| | Trauma | -0.99 | -1.6, -0.39 | 0.002 | **0.017** |
| **14,15-EPETrE** | Sepsis | — | — | | |
| | Aspiration | -0.03 | -0.48, 0.43 | >0.9 | >0.9 |
| | Other | 0.33 | -0.37, 1.0 | 0.4 | 0.5 |
| | Pneumonia | -0.07 | -0.54, 0.40 | 0.8 | 0.9 |
| | Trauma | -0.6 | -1.1, -0.05 | 0.032 | 0.11 |
| **14,15-EPETE** | Sepsis | — | — | | |
| | Aspiration | -0.64 | -0.99, -0.29 | <0.001 | **0.006** |
| | Other | -0.42 | -0.96, 0.11 | 0.12 | 0.3 |
| | Pneumonia | -0.43 | -0.79, -0.06 | 0.022 | **0.082** |
| | Trauma | -1.1 | -1.5, -0.64 | <0.001 | **<0.001** |
| **17,18-EPETE** | Sepsis | — | — | | |
| | Aspiration | -0.8 | -1.4, -0.18 | 0.012 | **0.054** |
| | Other | -0.24 | -1.2, 0.72 | 0.6 | 0.8 |
| | Pneumonia | -0.56 | -1.2, 0.09 | 0.088 | 0.2 |
| | Trauma | -1.5 | -2.2, -0.71 | <0.001 | **0.003** |
| **7,8-EPDPE** | Sepsis | — | — | | |
| | Aspiration | -0.54 | -1.0, -0.08 | 0.022 | 0.082 |
| | Other | 0.13 | -0.57, 0.84 | 0.7 | 0.9 |
| | Pneumonia | -0.02 | -0.50, 0.46 | >0.9 | >0.9 |
| | Trauma | -0.83 | -1.4, -0.28 | 0.003 | 0.024 |
| **13,14-EPDPE** | Sepsis | — | — | | |
| | Aspiration | -0.49 | -1.0, 0.06 | 0.079 | 0.2 |
| | Other | 0.07 | -0.77, 0.91 | 0.9 | >0.9 |
| | Pneumonia | -0.12 | -0.69, 0.45 | 0.7 | 0.8 |
| | Trauma | -0.62 | -1.3, 0.04 | 0.065 | 0.2 |
| **19,20-EPDPE** | Sepsis | — | — | | |
| | Aspiration | -0.28 | -0.83, 0.26 | 0.3 | 0.5 |
| | Other | 0.56 | -0.27, 1.4 | 0.2 | 0.4 |
| | Pneumonia | 0.04 | -0.53, 0.60 | 0.9 | >0.9 |
| | Trauma | -0.17 | -0.82, 0.48 | 0.6 | 0.8 |

| | | | | | |
|---|---|---|---|---|---|
| **9,10-diHOME** | Sepsis | — | — | | |
| | Aspiration | 1.7 | 0.88, 2.4 | <0.001 | **0.001** |
| | Other | 2.2 | 0.97, 3.4 | <0.001 | **0.007** |
| | Pneumonia | 0.54 | -0.26, 1.4 | 0.2 | 0.4 |
| | Trauma | 1.9 | 0.97, 2.8 | <0.001 | **0.002** |
| **12,13-diHOME** | Sepsis | — | — | | |
| | Aspiration | 1.5 | 0.69, 2.3 | <0.001 | **0.006** |
| | Other | 1.9 | 0.63, 3.1 | 0.003 | **0.025** |
| | Pneumonia | 0.31 | -0.53, 1.1 | 0.5 | 0.7 |
| | Trauma | 0.81 | -0.15, 1.8 | 0.1 | 0.2 |
| **5,6-diHETrE** | Sepsis | — | — | | |
| | Aspiration | -0.33 | -0.84, 0.19 | 0.2 | 0.4 |
| | Other | 0.33 | -0.46, 1.1 | 0.4 | 0.6 |
| | Pneumonia | 0 | -0.53, 0.54 | >0.9 | >0.9 |
| | Trauma | -0.92 | -1.5, -0.31 | 0.004 | **0.025** |
| **8,9-diHETrE** | Sepsis | — | — | | |
| | Aspiration | 0.03 | -0.46, 0.51 | >0.9 | >0.9 |
| | Other | 0.6 | -0.14, 1.3 | 0.11 | 0.3 |
| | Pneumonia | 0.01 | -0.49, 0.52 | >0.9 | >0.9 |
| | Trauma | -0.48 | -1.1, 0.09 | 0.1 | 0.2 |
| **11,12-diHETrE** | Sepsis | — | — | | |
| | Aspiration | -0.31 | -0.81, 0.18 | 0.2 | 0.4 |
| | Other | 0.73 | -0.03, 1.5 | 0.06 | 0.2 |
| | Pneumonia | 0.25 | -0.27, 0.76 | 0.3 | 0.5 |
| | Trauma | -0.77 | -1.4, -0.18 | 0.011 | **0.054** |
| **14,15-diHETrE** | Sepsis | — | — | | |
| | Aspiration | -0.11 | -0.61, 0.39 | 0.7 | 0.8 |
| | Other | 0.87 | 0.10, 1.6 | 0.027 | 0.1 |
| | Pneumonia | 0.12 | -0.40, 0.64 | 0.7 | 0.8 |
| | Trauma | -0.75 | -1.4, -0.15 | 0.014 | **0.064** |
| **19,20-diHDoPE** | Sepsis | — | — | | |
| | Aspiration | -0.23 | -0.81, 0.35 | 0.4 | 0.6 |
| | Other | 0.62 | -0.28, 1.5 | 0.2 | 0.4 |
| | Pneumonia | -0.16 | -0.77, 0.44 | 0.6 | 0.8 |
| | Trauma | -1 | -1.7, -0.32 | 0.005 | **0.029** |
| **9-oxo-ODE** | Sepsis | — | — | | |
| | Aspiration | 0.07 | -0.56, 0.69 | 0.8 | >0.9 |
| | Other | 0.13 | -0.84, 1.1 | 0.8 | >0.9 |
| | Pneumonia | 0.05 | -0.60, 0.71 | 0.9 | >0.9 |
| | Trauma | -1.1 | -1.8, -0.31 | 0.006 | **0.036** |
| **13-oxo-ODE** | Sepsis | — | — | | |
| | Aspiration | 0.64 | -0.01, 1.3 | 0.053 | 0.2 |
| | Other | 0.6 | -0.40, 1.6 | 0.2 | 0.4 |
| | Pneumonia | 0.36 | -0.32, 1.0 | 0.3 | 0.5 |
| | Trauma | -0.14 | -0.92, 0.64 | 0.7 | 0.9 |
| **9-oxo-OTrE** | Sepsis | — | — | | |
| | Aspiration | 0.42 | -0.31, 1.1 | 0.3 | 0.5 |
| | Other | 0.21 | -0.91, 1.3 | 0.7 | 0.9 |
| | Pneumonia | 0.17 | -0.58, 0.93 | 0.6 | 0.8 |
| | Trauma | -0.55 | -1.4, 0.32 | 0.2 | 0.4 |

| | | | | | |
|---|---|---|---|---|---|
| **15-oxo-EDE** | Sepsis | — | — | | |
| | Aspiration | -0.43 | -1.0, 0.16 | 0.2 | 0.3 |
| | Other | -0.06 | -0.97, 0.85 | 0.9 | >0.9 |
| | Pneumonia | 0.07 | -0.55, 0.69 | 0.8 | >0.9 |
| | Trauma | -1.1 | -1.8, -0.37 | 0.003 | **0.024** |
| **12-oxo-ETE** | Sepsis | — | — | | |
| | Aspiration | -0.17 | -0.56, 0.22 | 0.4 | 0.6 |
| | Other | -0.22 | -0.82, 0.38 | 0.5 | 0.6 |
| | Pneumonia | -0.04 | -0.45, 0.36 | 0.8 | >0.9 |
| | Trauma | -0.37 | -0.84, 0.09 | 0.12 | 0.3 |
| **15-oxo-ETE** | Sepsis | — | — | | |
| | Aspiration | 1 | 0.53, 1.5 | <0.001 | **0.001** |
| | Other | 0.45 | -0.28, 1.2 | 0.2 | 0.4 |
| | Pneumonia | 0.6 | 0.11, 1.1 | 0.018 | **0.07** |
| | Trauma | 0.1 | -0.47, 0.67 | 0.7 | 0.9 |
| **LXA4** | Sepsis | — | — | | |
| | Aspiration | 1 | 0.53, 1.5 | <0.001 | **0.001** |
| | Other | 0.45 | -0.28, 1.2 | 0.2 | 0.4 |
| | Pneumonia | 0.6 | 0.11, 1.1 | 0.018 | **0.07** |
| | Trauma | 0.1 | -0.47, 0.67 | 0.7 | 0.9 |
| **15-epi-LXA4** | Sepsis | — | — | | |
| | Aspiration | -0.18 | -0.58, 0.22 | 0.4 | 0.6 |
| | Other | -0.04 | -0.65, 0.57 | >0.9 | >0.9 |
| | Pneumonia | 0.09 | -0.33, 0.51 | 0.7 | 0.8 |
| | Trauma | -0.7 | -1.2, -0.23 | 0.004 | **0.028** |
| **LXA5** | Sepsis | — | — | | |
| | Aspiration | 0.47 | 0.19, 0.75 | 0.001 | **0.013** |
| | Other | 0.48 | 0.05, 0.91 | 0.027 | 0.1 |
| | Pneumonia | 0.2 | -0.09, 0.49 | 0.2 | 0.3 |
| | Trauma | 0.37 | 0.04, 0.70 | 0.03 | 0.1 |
| **LXB4** | Sepsis | — | — | | |
| | Aspiration | 0.77 | 0.14, 1.4 | 0.017 | **0.07** |
| | Other | 0.32 | -0.64, 1.3 | 0.5 | 0.7 |
| | Pneumonia | 0.62 | -0.03, 1.3 | 0.06 | 0.2 |
| | Trauma | 2.3 | 1.5, 3.0 | <0.001 | **<0.001** |
| **RvD1 + AT-RVD1** | Sepsis | — | — | | |
| | Aspiration | 0.58 | 0.10, 1.1 | 0.017 | **0.07** |
| | Other | 1 | 0.32, 1.8 | 0.006 | **0.033** |
| | Pneumonia | 0.28 | -0.22, 0.78 | 0.3 | 0.5 |
| | Trauma | 0.83 | 0.26, 1.4 | 0.005 | **0.029** |
| **RvD3** | Sepsis | — | — | | |
| | Aspiration | -0.04 | -0.37, 0.29 | 0.8 | >0.9 |
| | Other | -0.16 | -0.67, 0.34 | 0.5 | 0.7 |
| | Pneumonia | -0.23 | -0.58, 0.11 | 0.2 | 0.4 |
| | Trauma | -0.3 | -0.70, 0.09 | 0.13 | 0.3 |
| **AT-RvD3** | Sepsis | — | — | | |
| | Aspiration | 0.03 | -0.32, 0.38 | 0.9 | >0.9 |
| | Other | 0.04 | -0.49, 0.57 | 0.9 | >0.9 |
| | Pneumonia | 0.01 | -0.35, 0.38 | >0.9 | >0.9 |
| | Trauma | 0.06 | -0.35, 0.48 | 0.8 | 0.9 |

| | | | | | |
|---|---|---|---|---|---|
| **RvD4** | Sepsis | — | — | | |
| | Aspiration | 0.38 | 0.00, 0.76 | 0.05 | 0.2 |
| | Other | 0.82 | 0.24, 1.4 | 0.006 | **0.036** |
| | Pneumonia | 0.32 | -0.08, 0.71 | 0.11 | 0.3 |
| | Trauma | 0.57 | 0.12, 1.0 | 0.014 | **0.064** |
| **AT-RvD4** | Sepsis | — | — | | |
| | Aspiration | 0.11 | -0.45, 0.67 | 0.7 | 0.8 |
| | Other | 0.68 | -0.18, 1.6 | 0.12 | 0.3 |
| | Pneumonia | 0.13 | -0.46, 0.72 | 0.7 | 0.8 |
| | Trauma | 0.34 | -0.33, 1.0 | 0.3 | 0.5 |
| **RvD6** | Sepsis | — | — | | |
| | Aspiration | 0.08 | -0.35, 0.51 | 0.7 | 0.9 |
| | Other | 0.62 | -0.05, 1.3 | 0.068 | 0.2 |
| | Pneumonia | 0.19 | -0.26, 0.64 | 0.4 | 0.6 |
| | Trauma | 0.28 | -0.24, 0.79 | 0.3 | 0.5 |
| **AT-RvD6** | Sepsis | — | — | | |
| | Aspiration | 0.58 | 0.18, 0.98 | 0.005 | **0.031** |
| | Other | 0.47 | -0.14, 1.1 | 0.13 | 0.3 |
| | Pneumonia | 0.32 | -0.10, 0.74 | 0.13 | 0.3 |
| | Trauma | 0.77 | 0.29, 1.3 | 0.002 | **0.017** |
| **8-oxoRvD1** | Sepsis | — | — | | |
| | Aspiration | 0.45 | 0.06, 0.83 | 0.025 | **0.089** |
| | Other | 0.6 | 0.00, 1.2 | 0.048 | 0.2 |
| | Pneumonia | 0.31 | -0.09, 0.72 | 0.13 | 0.3 |
| | Trauma | 0.45 | -0.01, 0.92 | 0.056 | 0.2 |
| **RvD5 (n-3 DPA)** | Sepsis | — | — | | |
| | Aspiration | 0.49 | 0.18, 0.80 | 0.002 | **0.019** |
| | Other | 0.7 | 0.23, 1.2 | 0.004 | **0.028** |
| | Pneumonia | 0.36 | 0.04, 0.68 | 0.029 | 0.1 |
| | Trauma | 0.42 | 0.05, 0.79 | 0.026 | **0.095** |
| **RvE1** | Sepsis | — | — | | |
| | Aspiration | 1.2 | 0.34, 2.1 | 0.007 | **0.038** |
| | Other | 1.1 | -0.27, 2.5 | 0.11 | 0.3 |
| | Pneumonia | 0.12 | -0.80, 1.0 | 0.8 | >0.9 |
| | Trauma | 2.1 | 1.1, 3.2 | <0.001 | **0.003** |
| **RvE2** | Sepsis | — | — | | |
| | Aspiration | 0.06 | -0.32, 0.43 | 0.8 | 0.9 |
| | Other | 0.1 | -0.48, 0.67 | 0.7 | 0.9 |
| | Pneumonia | 0.27 | -0.12, 0.66 | 0.2 | 0.4 |
| | Trauma | -0.6 | -1.0, -0.15 | 0.009 | **0.045** |
| **22-OH-PD1** | Sepsis | — | — | | |
| | Aspiration | 0.66 | 0.18, 1.1 | 0.008 | **0.041** |
| | Other | 0.92 | 0.18, 1.7 | 0.016 | **0.067** |
| | Pneumonia | 0.63 | 0.13, 1.1 | 0.014 | **0.063** |
| | Trauma | 0.33 | -0.25, 0.90 | 0.3 | 0.5 |
| **Mar2** | Sepsis | — | — | | |
| | Aspiration | 0.24 | -0.20, 0.67 | 0.3 | 0.5 |
| | Other | 0.9 | 0.23, 1.6 | 0.009 | **0.045** |
| | Pneumonia | 0.33 | -0.12, 0.79 | 0.15 | 0.3 |
| | Trauma | -0.14 | -0.66, 0.38 | 0.6 | 0.8 |

| | | | | | |
|---|---|---|---|---|---|
| **Mar1 (n-3 DPA)** | Sepsis | — | — | | |
| | Aspiration | -0.31 | -0.83, 0.22 | 0.3 | 0.4 |
| | Other | 0.36 | -0.45, 1.2 | 0.4 | 0.6 |
| | Pneumonia | -0.26 | -0.81, 0.29 | 0.4 | 0.5 |
| | Trauma | -0.03 | -0.66, 0.60 | >0.9 | >0.9 |

Oxylipins were measured by LC/MS-MS. Tables shows beta (estimate) and confidence intervals for difference between sepsis-related ARDS and ARDS secondary to aspiration, other, pneumonia or trauma. with the Benjamini-Hochberg false discovery rate set for a significance threshold of $Q<0.1$. Significant results in **bold.** N=90

**Supplemental Table 5: Arachidonic Acid (AA) is negatively associated with proinflammatory cytokine IL-6, while n-3 DPA, AA and dihomo-Gamma-linolenic acid (DGLA) are negatively associated with both IL-6 and IL-8.**

| | IL-6 | | | | IL-8 | | | |
|---|---|---|---|---|---|---|---|---|
| | Beta | 95% CI | *P*-value | *Q*-value | Beta | 95% CI | *P*-value | *Q*-value |
| **ALA + GLA** | 0.03 | -0.07, 0.12 | 0.6 | 0.7 | 0.04 | -0.08, 0.16 | 0.5 | 0.6 |
| **SDA** | -0.1 | -0.19, 0.06 | 0.3 | 0.5 | -0 | -0.19, 0.12 | 0.6 | 0.7 |
| **EPA** | -0.1 | -0.18, 0.01 | 0.075 | 0.2 | -0.1 | -0.17, 0.07 | 0.4 | 0.5 |
| **n-3 DPA** | -0 | -0.09, 0.01 | 0.086 | 0.2 | -0.1 | -0.15, -0.02 | 0.009 | **0.03** |
| **DHA** | -0 | -0.10, 0.01 | 0.091 | 0.2 | -0.1 | -0.12, 0.01 | 0.085 | 0.13 |
| **LA** | 0 | -0.04, 0.04 | >0.9 | >0.9 | 0.04 | -0.01, 0.09 | 0.11 | 0.2 |
| **DGLA** | -0.1 | -0.14, 0.00 | 0.047 | 0.2 | -0.1 | -0.17, 0.00 | 0.045 | **0.085** |
| **AA** | -0.1 | -0.11, -0.03 | <0.001 | **0.014** | -0.1 | -0.12, -0.03 | 0.003 | **0.013** |
| **n-6 DPA** | -0 | -0.09, 0.01 | 0.12 | 0.2 | -0 | -0.09, 0.04 | 0.5 | 0.5 |

Lipidomics were measured by LC/MS-MS. Cytokines measured by multiplex ELISA. Table shows beta (estimate) and confidence intervals for fatty acids as compared to IL-6 and IL-8 with the Benjamini-Hochberg false discovery rate set for a significance threshold of $Q<0.1$. Significant results in **bold.** N=80

**Supplemental Table 6: Plasma normalized concentrations of oxylipins are associated with proinflammatory cytokines IL-6 and IL-8 in a cohort of patients with Acute Respiratory Distress Syndrome.**

| | IL-6 | | | | IL-8 | | | |
|---|---|---|---|---|---|---|---|---|
| | Beta | 95% CI | *P*-value | *Q*-value | Beta | 95% CI | *P*-value | *Q*-value |
| **13,14-dh-PGE** | -0.2 | -0.28, -0.07 | 0.001 | **0.047** | -0.4 | -0.48, -0.25 | <0.001 | **<0.001** |
| **13,14-dh-15k-PGE** | -0.1 | -0.20, -0.04 | 0.003 | **0.054** | -0.2 | -0.25, -0.05 | 0.003 | **0.008** |
| **PGE1** | -0.1 | -0.17, 0.02 | 0.12 | 0.3 | -0.3 | -0.35, -0.14 | <0.001 | **<0.001** |
| **15-keto-PGE1** | -0.1 | -0.20, -0.03 | 0.007 | **0.073** | -0.3 | -0.34, -0.15 | <0.001 | **<0.001** |
| **bicyclo-PGE1** | -0.1 | -0.17, 0.01 | 0.071 | 0.2 | -0 | -0.15, 0.07 | 0.4 | 0.5 |
| **2,3-dinor-PGE1** | -0.1 | -0.23, 0.06 | 0.2 | 0.4 | -0.1 | -0.27, 0.10 | 0.3 | 0.4 |
| **15-keto-PGE2** | -0.1 | -0.10, 0.00 | 0.058 | 0.2 | -0 | -0.10, 0.03 | 0.3 | 0.3 |
| **13,14-dh-15k-PGE2** | -0.1 | -0.15, 0.02 | 0.15 | 0.4 | -0.1 | -0.22, 0.00 | 0.042 | **0.067** |
| **bicyclo-PGE2** | -0 | -0.09, 0.03 | 0.4 | 0.6 | 0.02 | -0.05, 0.10 | 0.6 | 0.6 |
| **PGE3** | 0.29 | 0.11, 0.46 | 0.001 | **0.047** | 0.37 | 0.15, 0.59 | 0.001 | **0.004** |
| **PGJ2** | 0.01 | -0.04, 0.07 | 0.6 | 0.7 | 0.07 | 0.00, 0.13 | 0.045 | **0.07** |
| **d12-PGJ2** | -0 | -0.14, 0.10 | 0.7 | 0.8 | -0 | -0.19, 0.12 | 0.7 | 0.7 |
| **15d-d12,14-PGJ2** | -0.1 | -0.25, 0.10 | 0.4 | 0.6 | 0.02 | -0.20, 0.24 | 0.8 | 0.9 |
| **15d-d12,14-PGJ3** | -0 | -0.18, 0.15 | 0.8 | 0.9 | 0.02 | -0.19, 0.22 | 0.9 | >0.9 |
| **pgf1a** | -0 | -0.12, 0.03 | 0.2 | 0.4 | -0.1 | -0.16, 0.03 | 0.2 | 0.2 |
| **15-keto-PGF2a** | -0.1 | -0.20, -0.02 | 0.013 | 0.1 | -0.2 | -0.29, -0.08 | 0.001 | **0.004** |
| **19r(OH)PGF2a** | -0 | -0.29, 0.28 | >0.9 | >0.9 | 0.06 | -0.29, 0.42 | 0.7 | 0.8 |
| **8,isoPGF2a** | -0 | -0.09, 0.06 | 0.6 | 0.7 | -0.1 | -0.15, 0.04 | 0.2 | 0.3 |
| **6-keto-PGE1** | -0 | -0.16, 0.10 | 0.6 | 0.7 | -0.2 | -0.33, -0.02 | 0.032 | **0.054** |

| | | | | | | | | |
|---|---|---|---|---|---|---|---|---|
| **6,15-diketo-PGFa** | -0.1 | -0.15, 0.01 | 0.094 | 0.3 | -0.1 | -0.17, 0.04 | 0.3 | 0.3 |
| **2,3-dinor-TXB2** | -0 | -0.09, 0.03 | 0.3 | 0.5 | -0.1 | -0.17, -0.03 | 0.005 | **0.012** |
| **11,dh-2,3-dinor-TXB2** | -0.1 | -0.19, -0.04 | 0.002 | **0.047** | -0.2 | -0.25, -0.07 | <0.001 | **0.003** |
| **11-dh-TXB3** | -0.1 | -0.14, 0.03 | 0.2 | 0.4 | -0 | -0.11, 0.10 | >0.9 | >0.9 |
| **LTB4** | 0.05 | -0.03, 0.13 | 0.2 | 0.4 | 0.14 | 0.05, 0.24 | 0.004 | **0.009** |
| **LTB5** | 0.04 | -0.07, 0.14 | 0.5 | 0.6 | 0.19 | 0.07, 0.31 | 0.003 | **0.008** |
| **5(S),12(S)diHETE** | 0.06 | 0.00, 0.12 | 0.04 | 0.2 | 0.12 | 0.05, 0.19 | 0.001 | **0.004** |
| **5(S),15(S)diHETE** | -0 | -0.06, 0.03 | 0.5 | 0.7 | 0 | -0.06, 0.05 | 0.9 | >0.9 |
| **8(S),15(S)-diHETE** | 0.04 | -0.04, 0.13 | 0.3 | 0.5 | 0.06 | -0.05, 0.17 | 0.3 | 0.3 |
| **5(S),15(S)-diHEPE** | 0.09 | 0.00, 0.17 | 0.04 | 0.2 | 0.25 | 0.16, 0.34 | <0.001 | **<0.001** |
| **9-HODE** | 0.03 | -0.07, 0.13 | 0.6 | 0.7 | 0.12 | 0.00, 0.25 | 0.054 | **0.081** |
| **13-HODE** | 0.06 | -0.04, 0.17 | 0.2 | 0.4 | 0.17 | 0.04, 0.29 | 0.009 | **0.018** |
| **9-HOTrE** | 0.01 | -0.13, 0.15 | 0.9 | >0.9 | 0.15 | -0.03, 0.32 | 0.1 | 0.13 |
| **13-HOTrE** | 0.02 | -0.10, 0.13 | 0.8 | 0.8 | 0.17 | 0.02, 0.31 | 0.023 | **0.041** |
| **11-HEDE** | 0.11 | 0.01, 0.21 | 0.039 | 0.2 | 0.21 | 0.09, 0.34 | <0.001 | **0.004** |
| **15-HEDE** | 0.11 | 0.00, 0.23 | 0.055 | 0.2 | 0.21 | 0.07, 0.35 | 0.004 | **0.01** |
| **8-HETrE** | 0.09 | 0.01, 0.16 | 0.028 | 0.2 | 0.17 | 0.08, 0.27 | <0.001 | **0.002** |
| **5-HETrE** | 0.11 | 0.01, 0.21 | 0.038 | 0.2 | 0.25 | 0.13, 0.37 | <0.001 | **<0.001** |
| **5-HETE** | 0.04 | -0.04, 0.11 | 0.3 | 0.5 | 0.13 | 0.04, 0.23 | 0.004 | **0.01** |
| **8-HETE** | 0.06 | -0.01, 0.13 | 0.11 | 0.3 | 0.17 | 0.08, 0.25 | <0.001 | **0.001** |
| **9-HETE** | 0.05 | -0.07, 0.17 | 0.4 | 0.6 | 0.14 | -0.01, 0.29 | 0.062 | **0.091** |
| **11-HETE** | 0.06 | -0.03, 0.15 | 0.2 | 0.4 | 0.14 | 0.04, 0.25 | 0.009 | **0.018** |
| **12-HETE** | 0.01 | -0.13, 0.14 | >0.9 | >0.9 | 0.04 | -0.13, 0.21 | 0.7 | 0.7 |
| **15-HETE** | 0.07 | -0.01, 0.16 | 0.086 | 0.3 | 0.17 | 0.07, 0.27 | 0.001 | **0.004** |
| **20-HETE** | 0.05 | -0.03, 0.12 | 0.2 | 0.4 | 0.11 | 0.02, 0.20 | 0.016 | **0.03** |
| **tetranor-12-HETE** | 0.06 | 0.00, 0.13 | 0.059 | 0.2 | 0.11 | 0.03, 0.19 | 0.007 | **0.016** |
| **5-HEPE** | 0.03 | -0.06, 0.12 | 0.5 | 0.7 | 0.19 | 0.08, 0.29 | <0.001 | **0.004** |
| **8-HEPE** | 0.07 | -0.03, 0.18 | 0.2 | 0.4 | 0.25 | 0.13, 0.37 | <0.001 | **<0.001** |
| **9-HEPE** | 0.08 | -0.05, 0.20 | 0.2 | 0.4 | 0.28 | 0.14, 0.42 | <0.001 | **0.001** |
| **11-HEPE** | 0.04 | -0.07, 0.15 | 0.5 | 0.7 | 0.2 | 0.07, 0.34 | 0.003 | **0.008** |
| **12-HEPE** | -0 | -0.19, 0.14 | 0.7 | 0.8 | 0.07 | -0.14, 0.27 | 0.5 | 0.6 |
| **15-HEPE** | 0.08 | -0.03, 0.20 | 0.2 | 0.4 | 0.16 | 0.02, 0.30 | 0.028 | **0.05** |
| **18-HEPE** | 0.03 | -0.07, 0.14 | 0.5 | 0.7 | 0.12 | -0.01, 0.25 | 0.064 | **0.092** |
| **4-HDoHE** | 0.07 | -0.01, 0.14 | 0.073 | 0.2 | 0.14 | 0.05, 0.23 | 0.003 | **0.008** |
| **7-HDoHE** | 0.11 | 0.04, 0.19 | 0.005 | **0.061** | 0.18 | 0.09, 0.28 | <0.001 | **0.002** |
| **8-HDoHE** | 0.11 | 0.02, 0.21 | 0.018 | 0.12 | 0.21 | 0.09, 0.32 | <0.001 | **0.002** |
| **10-HDoHE** | 0.07 | -0.03, 0.17 | 0.2 | 0.4 | 0.19 | 0.06, 0.31 | 0.004 | **0.009** |
| **11-HDoHE** | 0.08 | -0.04, 0.19 | 0.2 | 0.4 | 0.18 | 0.04, 0.32 | 0.01 | **0.02** |
| **13-HDoHE** | 0.09 | -0.03, 0.20 | 0.2 | 0.4 | 0.18 | 0.04, 0.32 | 0.014 | **0.027** |
| **14-HDoHE** | 0 | -0.14, 0.15 | >0.9 | >0.9 | 0.1 | -0.08, 0.28 | 0.3 | 0.3 |
| **16-HDoHE** | 0.09 | -0.02, 0.19 | 0.1 | 0.3 | 0.21 | 0.08, 0.33 | 0.001 | **0.005** |

| | | | | | | | | |
|---|---|---|---|---|---|---|---|---|
| **17-HDoHE** | 0.08 | -0.02, 0.18 | 0.1 | 0.3 | 0.13 | 0.01, 0.25 | 0.039 | **0.064** |
| **20-HDoHE** | 0.19 | 0.07, 0.31 | 0.002 | **0.047** | 0.33 | 0.18, 0.47 | <0.001 | **<0.001** |
| **8,9-EPETrE** | 0.05 | -0.04, 0.13 | 0.3 | 0.4 | 0.15 | 0.05, 0.25 | 0.003 | **0.008** |
| **14,15-EPETrE** | 0.04 | -0.03, 0.11 | 0.3 | 0.4 | 0.08 | -0.01, 0.17 | 0.081 | 0.11 |
| **14,15-EPETE** | 0.03 | -0.03, 0.09 | 0.3 | 0.5 | 0.05 | -0.03, 0.12 | 0.2 | 0.3 |
| **17,18EPETE** | 0.03 | -0.07, 0.13 | 0.5 | 0.7 | 0.13 | 0.01, 0.25 | 0.032 | **0.054** |
| **7,8-EPDPE** | -0 | -0.10, 0.05 | 0.6 | 0.7 | 0.03 | -0.07, 0.12 | 0.6 | 0.6 |
| **13,14-EPDPE** | 0.08 | 0.00, 0.17 | 0.056 | 0.2 | 0.11 | 0.00, 0.22 | 0.045 | 0.07 |
| **19,20-EPDPE** | 0.12 | 0.04, 0.20 | 0.005 | **0.061** | 0.23 | 0.14, 0.33 | <0.001 | **<0.001** |
| **9,10-diHOME** | 0.08 | -0.06, 0.22 | 0.2 | 0.4 | 0.15 | -0.02, 0.31 | 0.092 | 0.13 |
| **12,13-diHOME** | 0.03 | -0.11, 0.16 | 0.7 | 0.8 | 0.14 | -0.03, 0.30 | 0.1 | 0.14 |
| **5,6-diHETrE** | 0.07 | -0.01, 0.15 | 0.11 | 0.3 | 0.16 | 0.06, 0.26 | 0.002 | **0.005** |
| **8,9-diHETrE** | 0.1 | 0.03, 0.18 | 0.009 | **0.08** | 0.2 | 0.12, 0.29 | <0.001 | **<0.001** |
| **11,12-diHETrE** | 0.03 | -0.05, 0.11 | 0.4 | 0.6 | 0.14 | 0.04, 0.24 | 0.006 | **0.013** |
| **14,15-diHETrE** | 0.07 | -0.01, 0.15 | 0.082 | 0.3 | 0.17 | 0.07, 0.27 | <0.001 | **0.004** |
| **19,20-diHDoPE** | 0.04 | -0.06, 0.13 | 0.4 | 0.6 | 0.02 | -0.09, 0.14 | 0.7 | 0.7 |
| **9-oxo-ODE** | 0.02 | -0.09, 0.12 | 0.7 | 0.8 | 0.08 | -0.05, 0.20 | 0.2 | 0.3 |
| **13-oxo-ODE** | 0.06 | -0.04, 0.16 | 0.2 | 0.4 | 0.17 | 0.05, 0.30 | 0.006 | **0.013** |
| **9-oxo-OTrE** | -0 | -0.14, 0.09 | 0.7 | 0.8 | 0 | -0.15, 0.14 | >0.9 | >0.9 |
| **15-oxo-EDE** | 0.04 | -0.05, 0.13 | 0.4 | 0.6 | 0.12 | 0.01, 0.23 | 0.035 | **0.058** |
| **12-oxo-ETE** | 0 | -0.06, 0.06 | >0.9 | >0.9 | -0 | -0.08, 0.07 | 0.9 | >0.9 |
| **15-oxo-ETE** | 0.02 | -0.06, 0.10 | 0.6 | 0.7 | 0.09 | 0.00, 0.19 | 0.062 | 0.091 |
| **LXA4** | 0.03 | -0.02, 0.09 | 0.3 | 0.4 | 0.08 | 0.02, 0.15 | 0.015 | **0.028** |
| **15-epi-LXA4** | 0.04 | -0.03, 0.10 | 0.3 | 0.4 | 0.1 | 0.02, 0.17 | 0.018 | **0.033** |
| **LXA5** | 0.04 | 0.00, 0.09 | 0.068 | 0.2 | 0.09 | 0.04, 0.15 | <0.001 | **0.004** |
| **LXB4** | -0.1 | -0.23, 0.00 | 0.048 | 0.2 | -0.2 | -0.35, -0.08 | 0.002 | **0.005** |
| **RvD1 + AT-RVD1** | 0.12 | 0.05, 0.20 | 0.002 | **0.047** | 0.18 | 0.09, 0.28 | <0.001 | **0.001** |
| **RvD3** | 0.02 | -0.03, 0.07 | 0.4 | 0.6 | 0.08 | 0.02, 0.14 | 0.011 | **0.021** |
| **AT-RvD3** | 0.06 | 0.01, 0.11 | 0.013 | 0.1 | 0.09 | 0.03, 0.15 | 0.002 | **0.006** |
| **RvD4** | 0.06 | 0.00, 0.12 | 0.04 | 0.2 | 0.12 | 0.05, 0.20 | 0.001 | **0.004** |
| **AT-RvD4** | 0.13 | 0.04, 0.21 | 0.004 | **0.061** | 0.23 | 0.13, 0.33 | <0.001 | **<0.001** |
| **RvD6** | 0.06 | -0.01, 0.12 | 0.11 | 0.3 | 0.18 | 0.11, 0.26 | <0.001 | **<0.001** |
| **AT-RvD6** | 0.05 | -0.02, 0.11 | 0.2 | 0.4 | 0.04 | -0.04, 0.12 | 0.3 | 0.4 |
| **8-oxoRvD1** | 0.02 | -0.05, 0.08 | 0.6 | 0.7 | 0.08 | 0.00, 0.16 | 0.041 | **0.065** |
| **RvD5** | 0.06 | 0.01, 0.11 | 0.012 | 0.1 | 0.11 | 0.05, 0.17 | <0.001 | **0.002** |
| **RvE1** | -0 | -0.17, 0.13 | 0.8 | 0.9 | 0.16 | -0.02, 0.35 | 0.081 | 0.11 |
| **RvE2** | 0.01 | -0.04, 0.07 | 0.6 | 0.7 | 0.04 | -0.03, 0.11 | 0.3 | 0.4 |
| **22(oh)-PD1** | 0.06 | -0.01, 0.14 | 0.1 | 0.3 | 0.17 | 0.07, 0.26 | <0.001 | **0.003** |
| **Mar1** | 0.07 | 0.01, 0.14 | 0.034 | 0.2 | 0.17 | 0.09, 0.25 | <0.001 | **<0.001** |
| **Mar1 (n-3 DPA)** | 0 | -0.08, 0.09 | >0.9 | >0.9 | 0.03 | -0.07, 0.14 | 0.5 | 0.6 |

Lipidomics conducted by LC/MS-MS. Cytokines measured by multiplex ELISA. Tables shows beta (estimate) and confidence intervales for fatty acids as compared to IL-6 and IL-8 with the Benjamini-Hochberg false discovery rate set for a significance threshold of *Q*<0.1. Significant results in **bold.** N=80
